# Visual uncertainty and task demands shape active sensing strategies in mice

**DOI:** 10.64898/2026.06.16.732525

**Authors:** Célia Benquet, Thomas Sainsbury, Léo Bruneau, Yang Lin, Chenchen Cai, Mariia Popova, Kayla Ponder, Lydia Ntanavara, Rachel Froebe, Zheng Tan, Paul Fahey, Katrin Franke, Luis M. Franco, Keaton Jones, Yizhou Chen, Reece Keller, Xaq Pitkow, Cristopher M. Niell, Andreas S. Tolias, Mackenzie Weygandt Mathis

**Affiliations:** EPFL Neuro, School of Life Sciences, EPFL, Genève, Switzerland; Department of Neuroscience, Center for Neuroscience and Artificial Intelligence, Baylor College of Medicine, Houston, TX,USA; Department of Ophthalmology, Byers Eye Institute, Stanford University School of Medicine, Stanford University, Palo Alto, CA, USA; Stanford Bio-X, Wu Tsai Neurosciences Institute, Stanford University, Stanford, CA, USA; Department of Biology and Institute of Neuroscience, University of Oregon, Eugene, OR, USA; Neuroscience Institute, Carnegie Mellon University, Pittsburgh, PA, USA; Robotics Institute, Carnegie Mellon University, Pittsburgh, PA, USA; Machine Learning Department, Carnegie Mellon University, Pittsburgh, PA, USA; Department of Electrical and Computer Engineering, Department of Computer Science, Rice University, Houston, TX, USA; NSF AI Institute for Artificial and Natural Intelligence (ARNI), Columbia Engineering School, New York, NY, USA; Department of Electrical Engineering, Stanford University, Stanford, CA, USA; Institute for Ophthalmic Research, Tübingen University, Tübingen, Germany

**Author notes:** These authors contributed equally (Célia Benquet, Thomas Sainsbury).

**Keywords:** active sensing, infotaxis, virtual reality, visual discrimination, decision making

## Abstract

In natural environments, animals actively sample visual information to guide behavior. Sensory feedback is dynamic and often requires active movements, whether saccading across the lines of this page or walking through a park. From high-acuity vision in hawks to low-acuity mice, many animals actively navigate to seek information, which can be called infotaxis. Although mice have relatively low-acuity vision, they still rely on sight for critical behaviors including navigation and prey capture. Yet, how sensitive they are to visual information and performing infotaxis has not been established. Here, we develop a virtual reality object discrimination task to investigate visual decision-making under naturalistic conditions. We show that mice perform infotaxis by actively seeking out informative views to guide their choices. Stimulus manipulations confirm that this strategy is modulated by the amount of available visual information. These results reveal that mice use principled active strategies to resolve visual uncertainty, highlighting a key role for information-seeking in natural vision.

**Highlights:**

- Mice perform active visual sensing in a free-range VR object discrimination task.
- They perform infotaxis and select the correct object under variable visibility.
- Decisions decoded from head-body movements and speed reflect occlusion sensitivity.

## Introduction

Traditional vision studies have largely assumed a passive observer (1–6), yet animals in natural environments actively gather visual information to guide behavior (7, 8). This process, called active sensing, involves the use of movement to selectively sample sensory input in a goal-directed manner (9–11) From a computational normative perspective, active sensing forms a closed loop between an observer, which infers task-relevant variables from incoming sensory data, and a planner, which selects actions that yield informative observations (12). This framework connects sensory processing and motor behavior through continuous, goal-directed information gathering (13– 15).

The implementation of active sensing varies dramatically across species depending on sensory anatomy, leading to distinct sampling strategies. Species with a high-resolution portion of the retina, such as a fovea or area centralis, benefit from rapidly shifting that high-acuity region via eye movements (16). In primates, this is characterized by rhythmic sampling of the visual scene where saccadic eye movements and stereotyped fixation durations aligns the fovea with task-relevant visual features (9, 17–22). By contrast, species such as mice with more uniform retinal resolution (23, 24) rely more heavily on head and full body movements to leverage cues such as motion and position parallax, while their eye movements occur mostly in tandem with head movements to compensate for reorientation (25–27). Despite these anatomical differences, mice still depend on vision for a range of ecologically relevant behaviors including threat detection (28, 29), foraging (30, 31), spatial navigation (32, 33), and depth perception (27, 34). However, no paradigm has yet been developed to systematically measure specific active visual infotaxis strategies in this species (11).

To address this gap, we developed a free-range psychophysics paradigm for laboratory mice to test the hypothesis that mice engage in active visual sensing based on task demands. We designed a virtual reality-based object discrimination task that used real-time pose estimation to control visual stimuli, enabling dynamic manipulation of visual occlusion via a wall-like aperture that introduced depth parallax. Specifically, changes in viewing angle exposed different parts of the object from behind occluders: by moving closer to the screen, mice could reveal more of the occluded object. Although head-fixed virtual reality approaches enable precise visual control at the cost of natural movements (35– 37), our freely moving paradigm allowed mice to actively control their viewing angle through body and head movements, enabling them to actively sample visual information while we maintained complete knowledge of the visual stimuli. We found that mice modulated their behavior based on the amount of available information, consistent with an infotaxis strategy that seeks to maximize information gain (7, 38– 40). Trial-by-trial decisions could be decoded from head-body movements and locomotor speed, with decoding accuracy varying based on object visibility. This indicated that mice adapted their sensing strategies in response to visual uncertainty. These findings demonstrate that mice perform active visual sensing and flexibly deploy sensorimotor strategies to resolve object ambiguity under variable visibility conditions.

## Results

### An active sensing task for mice

To investigate whether mice engage in active sensing during visual decision-making, we developed a freely moving virtual reality (VR) system that provides real-time, closed-loop visual feedback from the point of view of the animal, based on its position and posture (Figure 1**A**, STAR Methods). We used an off-axis projection of the scene (Figure 1**B**) so that visual scenes were rendered dynamically from the instantaneous viewpoint of the animal and the correct motion parallax was preserved. Nearer virtual objects shifted more with head movements than distant ones, producing a realistic three-dimensional visual experience. To generate the scene from the animal’s point of view, we integrated *DeepLabCut-Live!* for real-time pose estimation (41)

(Figure S1**A:C**) with Unity for instantaneous viewpoint rendering (Figure S1**D**), managed via a custom Python-based bridge (Figure 1**C**). This ensured low-latency, closed-loop feedback (mean ± s.e.m. per lab: Tolias lab: 80.62 ± 2.40 ms, n = 5; MLAI: 56.10 ± 0.83 ms, n = 6; Niell lab: 59.70 ± 0.40 ms, n = 5; Figure 1**D**, Figure S1**E:G**). We also controlled for acoustic stability of the experimental environment and showed that the system does not emit task-aligned noise that could help the mice in any way (Figure S1**H**). In brief, this system enabled naturalistic interaction with visual stimuli while maintaining precise experimental control, allowing us to design tasks that probe visual information-gathering behavior. We implemented identical setups and protocols across three laboratories, establishing a reproducible framework for studying active vision.

**Figure 1.**
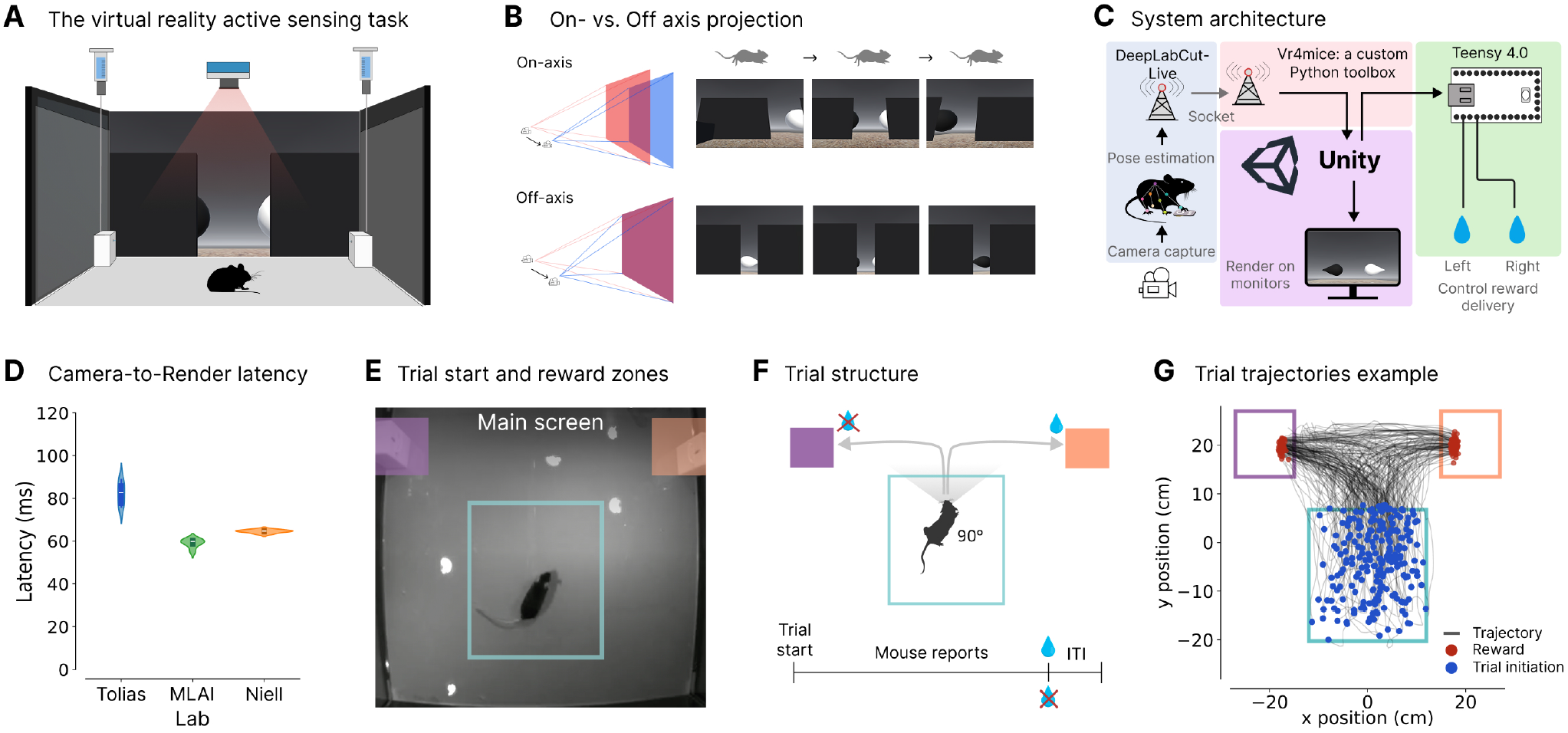
A closed-loop virtual-reality active sensing task for mice. **A**: Schematic of the virtual reality arena. The mouse moves freely around while a virtual world renders dynamically on the front screen. Lick ports are situated at the front sides. A high-speed overhead camera acquires images at 100Hz. **B**: On-vs. Off-axis projection: Above, On-axis projection: camera viewing angle moves out of projection plane. When the mouse moves right, the camera frustum remains symmetric relative to the center, leading to inaccurate motion parallax. Below, Off-axis projection: camera viewing angle is fixed to the projection plane. When the mouse moves right, the projection matrix adjusts to its viewpoint, accurately aligning virtual content to its perspective. **C**: Virtual reality software architecture. DeepLabCut-Live captures images for real-time pose estimation. Head x/y positions and running direction are extracted via our custom Python toolbox, `vr4mice`, and sent to Unity for rendering. `vr4mice` also controls water reward delivery via a Teensy *±* 4.0 for correct reports. **D**: Total camera-to-render latency across labs, summing DeepLabCut-Live and Unity latencies (Figure S1**E:G**). Mean s.e.m.: 80.62 ± 2.40 ms (Tolias; n=5), 59.70 ± 0.40 ms (Niell; n=5) and 56.10 ± 0.83 ms (MLAI; n=6). **E**: Arena top view overlaid with the trial-start area (blue) and report areas (orange: right; purple: left). Areas not visible to the mouse. Main screen at the front. **F**: Trial structure. A trial starts when the mouse enters the initiation area, aligns its running direction with the screen, and meets a velocity threshold. The mouse reports the object of interest location by entering the corresponding side reward port. **G**: Example trial trajectories from a trained mouse during discrimination (session: Oribi, 2024-08-14). Blue/Red dots: start/end positions. See also Figure S1.

To initiate a trial, the mouse had to enter a trial initiation zone (non-visible to the mouse) in the arena while decreasing its velocity and facing the front screen for 250 ms (blue rectangle in Figure 1**E**). This caused two objects to appear on the screen in front of the mouse. It could then report the location of the target object by going to the right or left lick port (purple and orange shaded rectangles in Figure 1**E**). If the mouse reported correctly by going to the lick port on the same side as the target object, it received a water reward (3–5 *µ*L), and the objects disappeared until the mouse initiated another trial (inter-trial interval). If the mouse reported incorrectly, no water reward was delivered and the inter-trial interval was triggered directly (Figure 1**F**). Across the three laboratories, on the majority of trials mice displayed trajectories that formed a J-shape as they moved toward the lick ports (Figure 1**G**).

Mice were trained in a two-step regimen (Figure 2**A**, Figure S2**A**): first an object detection task, followed by an object discrimination task. In the detection phase, mice learned to report the correct location of an object of interest (a white teardrop). The object is stationary in the virtual coordinate system, but the rendered viewpoint is coupled to the mouse’s movements so that the available visual information at any given point in space depends on the mouse’s position. To facilitate learning trial initiation, the detection phase was split into two subtasks. First, the mice had to enter the start zone and remain for 100 ms to initiate a trial. Once they reached sufficient performance (above a 70% reward rate on more than 125 trials within an hour, see STAR Methods), they were switched to the standard trial-initiation criteria used for the rest of the paradigm: entering the start zone for a longer duration of 250 ms. After reaching criterion on this as well, they graduated to the discrimination task, where the target object and a distractor object (black teardrop) were presented simultaneously, and mice had to report the target’s location.

**Figure 2.**
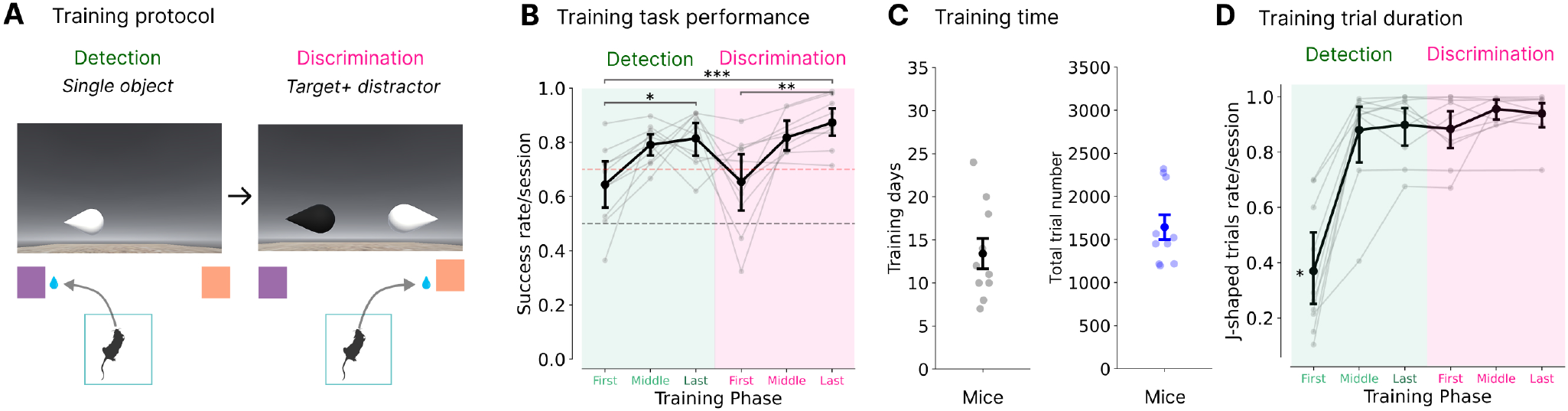
Mice can learn to discriminate objects based on luminance in a motion-parallax correct virtual reality world. **A**: Mice first learned to detect a single white teardrop (detection phase). Next, a black teardrop distractor was added (discrimination phase); mice still reported the white teardrop’s position. **B**: Success rate across training phases. Green: detection phase, showing first/last sessions without velocity threshold (First/Middle), and the last session with velocity threshold (Last). Pink: first, middle (if applicable), and last sessions of the discrimination phase (with distractor). One-way ANOVA, significant training stage effect: F(1, 53) = 8.13, *p* = 0.006. Tukey HSD tests, significant pairwise differences, : *∗ ∗ ∗ p* = 0.001, : *∗∗ p* = 0.002, : *∗ p* = 0.025 (n=10), Figure S2**B. C**: Averaged training days (Left) and corresponding total number of trials (Right) until mice were considered experts at the discrimination task. Mean *±* s.e.m.: 13.4 ± 1.76 days, 1644.0 *±* 143.6 trials (n=10). **D**: Mean J-shaped trials rate across training. One-way ANOVA, significant training stage effect: F(1, 53) = 29.53, *p <* 0.001. : *∗* Tukey HSD tests, significant pairwise differences between first detection session and others, *p <* 0.001 (n=10), Figure S2**D**. See also Figure S2.

Throughout training, mice showed an increase in performance, with a significant improvement occurring between early and late training sessions for the detection and discrimination phases (one-way ANOVA significant effect of training stage, F(1, 53) = 8.13, *p* = 0.006; Tukey HSD, First vs. Last detection *p* = 0.025, First vs. Last discrimination *p* = 0.002, First detection vs. Last discrimination *p* = 0.001; Figure 2**B**, Figure S2**B**). Mice reached expert performance (same criterion as for detection phase: 70% success rate on at least 125 trials within 60 minutes, see STAR Methods; number of trials per session significantly higher than the first training session for all subsequent sessions; Tukey HSD *p <* 0.001; Figure S2**C**) in 13.4 ± 1.76 days for a total of 1644 ± 143.6 trials on average (mean ± s.e.m.; Figure 2**C**). Following the first detection session, J-shaped trials became significantly more frequent (Tukey HSD, *p <* 0.0001; Figure 2**D**, Figure S2**D**), while trial tortuosity decreased significantly (Figure S2**E**; Tukey HSD *p <* 0.0001, see STAR Methods). Simultaneously, reporting time dropped significantly (Tukey HSD, First vs. Middle detection *p* = 0.009, overall *p <* 0.025; Figure S2**F**), while a significant choice bias emerged transiently upon presentation of the distractor (one-sample t-test, First discrimination: t = 3.03, *p* = 0.014; Middle discrimination: t = 5.20, *p* = 0.007, all others *p >* 0.25; Figure S2**G**). Overall, these results confirm that mice reliably learned a consistent set of actions consisting of J-shaped trajectories to solve the task.

### Active sensing under partial visibility of objects

To investigate whether mice could perform visual active sensing, we introduced a task where partial occlusion of the objects occurred by presenting them behind wall-like occluders of randomly-sampled size (99% versus 10% of objects visible when viewing from the center of the trial initiation area, Figure 3**A**; equal frequency aperture conditions: paired t-test, t(42) = 0.49, *p* = 0.63, Figure S3**A**; Video S1). We hypothesized that this manipulation would increase task demands and require mice to modulate their behavioral strategy accordingly. Specifically, if mice perform active sensing, they should adjust their information-seeking behavior in response to reduced object visibility, upon the narrow-aperture condition, as less visual information would initially be accessible through the aperture.

**Figure 3.**
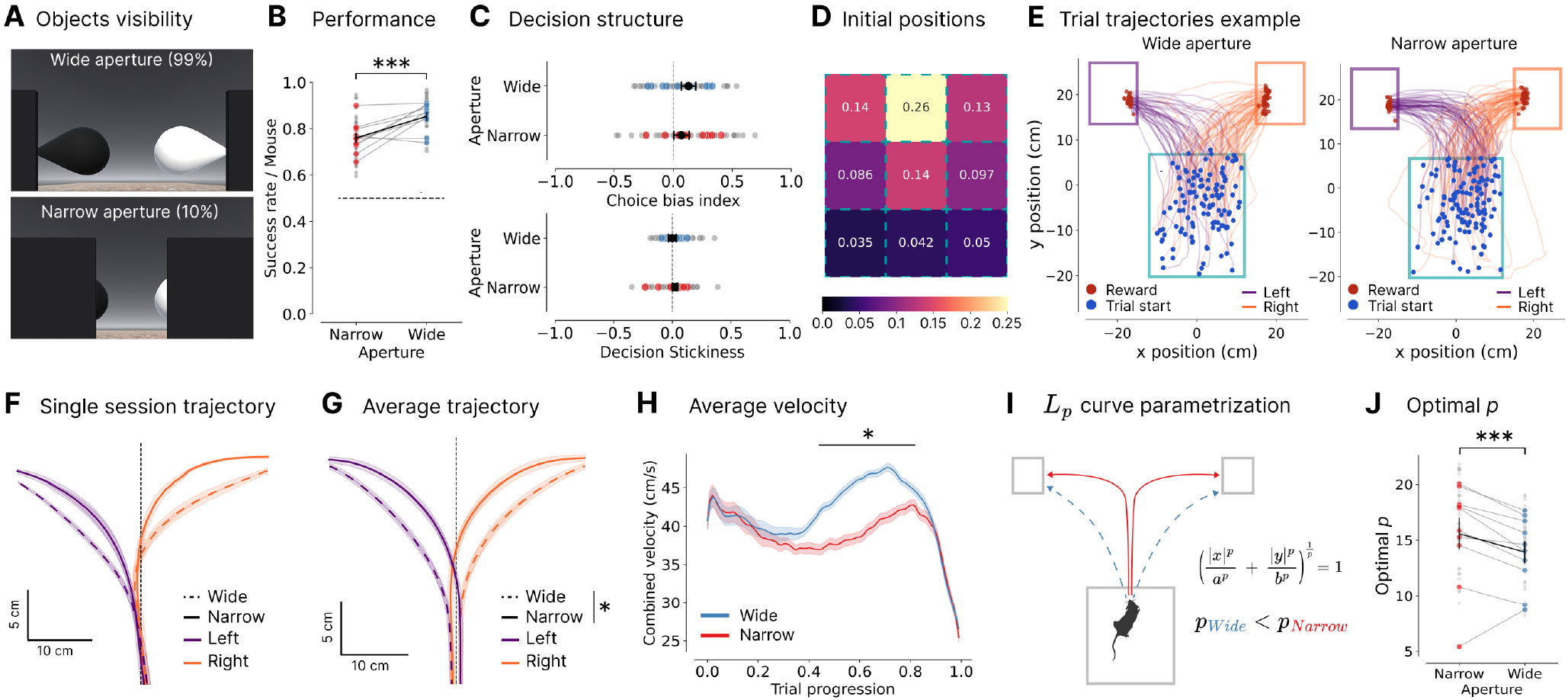
Mice stay more central, move slower and orient themselves more toward the objects as they approach the screen when faced with obstructed objects. **A**: Extent of the visual aperture during the dual-occlusion task: wide (99% of the objects visible) vs. narrow (10% of the objects visible). **B**: Success rate per aperture. Mean *±* s.e.m.: 0.76 *±* 0.01 (narrow) and 0.86 *±* 0.01 (wide). Paired t-test: t(42) = 6.98, *p* = 1.55 *×* 10^*−*8^. **C**: Influence of previous trial outcome on current choice. Above: Side choice bias per aperture. 0 = no bias, *±* 1 = full Left/Right. Mean s.e.m.: *±* 0.158 0.042 *±* (narrow) and 0.052 *±*0.035 (wide). One-sample t-test vs. no bias across mice: narrow, t = 2.09, *p* = 0.066; wide: t = 1.07, *p* = 0.311. Below: Decision stickiness per aperture. 0 = no stickiness to the previous decision, +1 = repeat, −1 = switch. Mean *±* s.e.m.: 0.025 *±* 0.021 (narrow) and 0.033 *±* 0.018 (wide). One-sample t-test vs. no stickiness across mice: narrow, t = 0.01, *p* = 0.995, wide, t = 1.08, *p* = 0.307. **D**: 3 x 3 initial position distribution across trials (n=10 mice, s=43 sessions). See Figure S3**E** for choice-specific distributions. **E**: Example trial trajectories for wide (left) and narrow (right) apertures (session: Pheasant, 2024-08-15). Mean trajectory *±* s.e.m. (shaded) per aperture and choice for **F**: individual example mouse (Pheasant, s=5 sessions; Figure S3**F**), and **G**: mean across mice (n=10 mice, s=43 sessions). Two-way repeated-measures ANOVA, significant effects of aperture: F(1, 42) = 68.65, spatial bin: F(40, 1680) = 89.21, and interaction: F(40, 1680) = 6.41; all *p <* 0.0001. *∗* : FDR-corrected paired t-test with multiple comparisons across y bins (0-18 cm in the arena), significant aperture effect (*p <* 0.05) from 0 cm onward; Figure S3**G**.Trajectories shown from arena midpoint (y-axis) to screen; dashed line indicates x-axis center. **H**: Mean velocity *±* s.e.m. per aperture (n=43). Two-way repeated-measures ANOVA, significant effects of aperture: F(1, 42) = 58.36, trial progression: F(99, 4158) = 48.18, and interaction: F(99, 4158) = 35.61; all *p <* 0.0001. *∗* : FDR-corrected paired t-test with multiple comparisons across trial progression, significant aperture effect (*p <* 0.05) on the 40-80% portion of the trial, Figure S3**H**. **I**: Depending on the mouse’s forward-movement toward the screen, we can approximate a trajectory using a constant L_*p*_ norm curve, parameterized by *p* and the horizontal and vertical length scales *a* and *b*. Larger *p* indicates sharper curvature, reflecting further forward movement before committing to a decision under narrow occlusion. **J**: Optimal *p* parameter from single-trial L_*p*_ curve fits. Mean *±*s.e.m.: 16.68*±* 0.58 (narrow) and 14.53 *±* 0.49 (wide). *∗ ∗ ∗* : paired t-test, t(42) = *−* 7.12, *p* = 9.78 *×* 10^*−*9^. For **B, C** and **J**: Black: mean *±* s.e.m.; colored: per-mouse; grey: per-session. See also Figure S3 and Video S1.

Mice maintained high discrimination accuracy in both occluded and unoccluded trials (mean narrow = 0.76 ± 0.01; mean wide = 0.86 ± 0.01; Figure 3**B**, Figure S3**B**). The significant decrease in accuracy during occlusion (paired t-test: t(42) = 6.98, *p* = 1.55 × 10^*−*8^) effectively highlights the increased sensory complexity posed by hidden objects while confirming the mice’s ability to solve more demanding conditions. They localized the object positions without a population-level side bias (one-sample t-test vs. no bias: narrow, t = 2.09, *p*=0.066; wide: t = 1.07, *p* = 0.311; mean narrow: 0.129 ± 0.062, mean wide = 0.069 ± 0.064), or decision stickiness (one-sample t-test vs. no stickiness to the previous choice; narrow: t = 0.01, *p* = 0.995, wide: t = 1.08, *p* = 0.307) at both aperture levels, even if subtle individual biases existed (Figure 3**C**; Figure S3**C** for results per laboratory). High rates of J-shaped trials also indicate strong task engagement (mean: 0.972 ± 0.0044; Figure S3**D**). Furthermore, we found no difference in the trial initial position between choice sides (paired t-test: FDR-corrected p-value, *p >* 0.05 for all spatial bins; Figure 3**D**, Figure S3**E**). This suggests that mice naturally move toward a specific position before task initiation, resulting in a consistent starting view of the objects regardless of the subsequent decision. This consistent engagement and lack of systematic bias – even under occlusion – sets the stage for examining underlying information-seeking strategies.

A behavioral strategy that would enable mice to perform the task accurately when faced with occlusion is to move closer to the screen while remaining near the arena center. This would result in a wider viewing angle through the aperture, allowing the mice to observe more surface area of the objects of interest. We observed such a pattern across trials (Figure 3**E**) and in the per-mouse averaged trajectories (Figure 3**F**, Figure S3**F**). As they moved toward the front screen, all mice stayed in the central portion of the arena more in the narrow than in the wide aperture condition. Overall, we observed a significant difference between averaged trajectories per aperture condition, consistent with our initial hypothesis (two-way repeated-measures ANOVA significant effect of aperture: F(1, 42) = 68.65, spatial bin: F(40, 1680) = 89.21, and interaction: F(40, 1680) = 6.41; all *p <* 0.0001; FDR-corrected paired t-test with multiple comparison across y bins: *p <* 0.05; Figure 3**G**, Figure S3**G**).

In addition, if movements were independent of the information available, both conditions should exhibit identical velocity profiles, and if mice were unsure about their decision, this should be reflected in their velocity. Analysis of the animals’ velocity revealed a significantly slower acceleration between 40% and 80% of trial progression in the narrow aperture condition compared to the wide aperture trials (two-way repeated-measures ANOVA significant effects of aperture: F(1, 42) = 58.36, time progression: F(99, 4158) = 48.18, and interaction: F(99, 4158) = 35.61; all *p <* 0.0001; FDR-corrected paired t-test with multiple comparison across y bins: *p <* 0.05; Figure 3**H**, Figure S3**H**). When examining the velocity components independently, the y-velocity (forward movements, toward the screen; Figure S3**I**) was lower throughout the trial under occlusion, while the x-velocity (lateral movements, toward the reward ports; Figure S3**J**) showed a delayed peak in the narrow condition. This indicates that the increased sensory demand of the narrow aperture led to a more cautious velocity profile characterized by a slower approach to the screen and a delayed commitment to a choice side, likely to maximize the time available for information gathering.

Beyond velocity, the spatial orientation of the mice also diverged according to the available information. The running direction differed significantly between conditions from trial onset, except between 70% and 80%, which aligns with the observed velocity decrease in the narrow condition (two-way repeated-measures ANOVA, significant effects of aperture: F(1, 42) = 19.06, time progression: F(99, 4158) = 1344.43, and interaction: F(99, 4158) = 33.54; all *p <* 0.0001; FDR-corrected paired t-test with multiple comparisons: *p <* 0.05; Figure S3**K-L**). This suggests that mice adapted their behavioral trajectory from the very beginning of the trial based on the presented level of occlusion. Moreover, the angle of the head relative to the body showed significant differences from 60% to 90% of the trial (two-way repeated-measures ANOVA, significant main effects of aperture: F(1, 42) = 30.96, time progression: F(99, 4158) = 92.55, and interaction: F(99, 4158) = 5.83; all *p <* 0.0001; FDR-corrected paired t-test with multiple comparisons: *p <* 0.05; Figure S3**M-N**). This postural shift in the later stage is consistent with the sudden turn towards the reward port observed in the narrow condition’s average trajectory, suggesting a rapid commitment once sufficient information has been integrated.

Taken together, these results suggest that when faced with a high degree of occlusion (narrower aperture), mice moved closer toward the screen for longer in the trial before making a decision. One possible reason for this is that mice move to a position within the arena where they increase their visual angle through the aperture, allowing them to better view the objects of interest to then correctly classify their location. As such, these trials can be thought of as having two distinct phases: a *search phase* where mice seek out object information that is hidden by the occlusion, followed by a *decision phase* where – once enough information has been integrated (i.e., once they are confident enough) – they commit to going to one of the reward ports. In the wide-aperture trials, on the other hand, mice can readily obtain this information from further back in the arena and can commit to a decision without entering a search phase, or at least with only a shortened one.

To better quantify this, we fitted the individual experimental trial movement trajectories to L_*p*_ curves by finding the optimal *p* for each trial (STAR Methods). The L_*p*_ family of curves provides a flexible way to capture trajectory curvature, where larger *p* values correspond to more sharply curved (i.e., more J-shaped) paths, consistent with the intuition that mice exhibit more pronounced J-shaped trajectories in narrow aperture trials (Figure 3**I**). We observed a significant difference in the average optimal *p* between aperture conditions (mean narrow = 16.68 ± 0.58, mean wide = 14.53 ± 0.49; paired t-test across sessions t(42) = − 7.12, *p* = 9.77 × 10^*−*9^; Figure 3**J**, Figure S3**O**), indicating sharper turns when objects are occluded. Similarly, the significantly higher tortuosity observed in narrow aperture trials compared to wide ones (mean narrow = 1.56 ± 0.03, mean wide = 1.33 ± 0.01, paired t-test: t(42) = −6.85, *p* =− 2.41 × 10^*−*8^; Figure S3**P**) indicated that mice shifted to more complex search paths when sensory information is constrained. Analysis of the most tortuous forward paths (top 90^th^ percentile) revealed that these were predominantly narrow-aperture trials (83.6%; Figure S3**Q**). These trajectories sometimes showed “change of mind” patterns, characterized by an initial movement toward one target followed by a correcting redirection toward the other. Such behavior indicated that the trajectories were not ballistic or pre-planned. Instead, they suggested that mice actively adjusted their behavior in reaction to late-arriving evidence, utilizing incoming information to update an internal decision variable and self-correct mid-trial (42, 43).

We also trained a batch of mice (n=3) on the inverse contrast experiment (black target, white distractor) and observed similar behavioral changes (Success rate: narrow: 0.74 ± 0.02, wide: 0.90 ± 0.02; t(14) = 4.35, *p* = 6.67 × 10^*−*4^; Trajectory: two-way repeated-measures ANOVA, significant effects for aperture: F(1, 14) = 11.96, *p* = 0.0038, spatial bin: F(14, 196) = 221.97, *p <* 0.0001 and interaction: F(14, 196) = 5.80, *p <* 0.0001; FDR-corrected paired t-test with multiple comparisons across y bins, significant aperture effect from 2.5 cm of the initiation area top onward; Combined velocity (m/s): two-way repeated-measures ANOVA, significant effect of aperture: F(1, 14) = 63.07, time bin: F(99, 1386) = 46.39, and interaction: F(99, 1386) = 22.22; all *p <* 0.0001; paired t-test with multiple comparisons across the trial progression, significant aperture effect at 40-80% trial; Decision points distance to the screen (cm): narrow: 14.64 ± 0.67, wide: 18.29 ± 0.52; t(14) = 3.76, *p* = 0.021; STAR Methods;

Figure S3**R**). This suggests that the animals’ navigation and sensing strategies are driven by object identity and task rules rather than a luminance-based detection of the white object or confusion of the black object with the arena walls.

### A dynamic choice variable predicts infotaxis behavior

We hypothesized that aperture width should shift the position at which mice commit to a choice: in narrow-aperture trials, decisions should occur closer to the screen than in wide-aperture trials. To resolve the time course of infotaxis, we trained a time-resolved multi-model framework consisting of ten independent logistic regression models – one per 10% window of the trial – to predict the animal’s upcoming choice (left versus right reward port). One model was trained per time segment and session in a leave-one-session-out procedure (n=43; Figure 4**A**; STAR Methods). This allowed us to estimate when behavior first became predictive of the final decision. To determine which features best capture the decision process, we compared ten candidate feature sets using the Bayesian Information Criterion (BIC; STAR Methods). The best-performing model included only kinematic and orientation features related to lateral displacement (Figure S4**A**; Figure S4**C:F** for regressor time courses). Thus, lateral movements provide the most informative behavioral signature of choice formation. This is consistent with the task geometry, where the x-axis defines the left–right decision axis in the arena. Analysis of regression weights along the time segments revealed a three-stage evolution in the relative importance of these features (Figure 4**B**, Figure 4**G**): in the beginning of the trial (0-30%), predictive power is negligible; during the mid-trial deliberating phase (40-80%), x-velocity and orientation emerge as the primary predictors; finally, as the mice approach the choice (80-100%), x-position becomes the dominant feature.

We used the time-specific probability estimates of these models as a dynamic choice variable (43, 44) along trial progression (Figure 4**C-D**). First, we observed that the prediction accuracy of the model throughout the trial per occlusion condition showed a significant shift between conditions from 30% to 90% (two-way repeated-measures ANOVA significant effects of aperture: F(1, 42) = 26.19, trial progression: F(49, 2058) = 794.25, and interaction: (F(49, 2058) = 9.63; all *p <* 0.0001; FDR-corrected *p <* 0.05; Figure 4**E**, Figure S4**H**). Thus, the model reached the same level of confidence significantly later in the trial under narrow apertures. This indicates that the behavioral features used as model regressors varied sufficiently between aperture conditions to drive this temporal predictability difference, suggesting distinct behavioral strategies.

To quantify the timeline of the choice process, we defined the decision point – i.e., the transition from information gathering to directed navigation – as the moment the model confidence reached a specific certainty threshold (Figure 4**F-G**). To determine the threshold that most accurately reflected the mice turning dynamics, we evaluated certainty levels ranging from 50% (random) to 90%. A threshold of 80% was found to best capture the onset of the animal’s lateral trajectory toward the reward port (Figure S4**B**; STAR Methods). Decision points defined as such occurred significantly closer to the screen in the narrow aperture condition compared to its wide counterpart (means ± s.e.m. narrow: 13.79 ± 0.44 cm and wide: 17.54 ± 0.42 cm, mean difference = 3.75. ∗ ∗ ∗ : paired t-test, t(42) = 6.03, *p* = 3.56 × 10^*−*7^; Figure 4**H**, Figure S4**I**).

Overall, as task demands increased – with the narrow aperture increasing visual uncertainty and making the task more challenging to perform as evidenced by their drop in performance – mice delayed their decision, spending more of the trial getting closer to the front screen. This resulted in them moving to more informative viewpoints (i.e., more of the objects was visible; information gain between initial and decision point positions, paired t-test, t(42) = − 10.97, *p* = 6.67 × 10^*−*14^; Figure S4**J**). We then analyzed how movement dynamics evolved as evidence accumulated during the search phase. Specifically, we examined the velocity toward the screen (y-axis) as a function of the information content aligned to the decision. In both conditions, velocity increased rapidly until approximately 0.4 seconds before the decision point. Their trajectories then diverged: in narrow-aperture trials, velocity peaks at 0.22 seconds before the decision before decreasing (*r* = − 0.68, *p <* 0.001), while in wide-aperture trials, velocity peaks at 0.20 seconds before the decision but stays overall high (*r* = 0.42, *p* = 0.007; Figure 4**I**). Then both conditions showed an abrupt deceleration immediately preceding the decision, corresponding to the physical turn to the reward port. This demonstrates evidence-driven modulation of velocity, reflecting reactive behavioral adjustments to task difficulty and visual uncertainty and consistent with an extended infotaxis strategy when faced with object occlusion.

**Figure 4.**
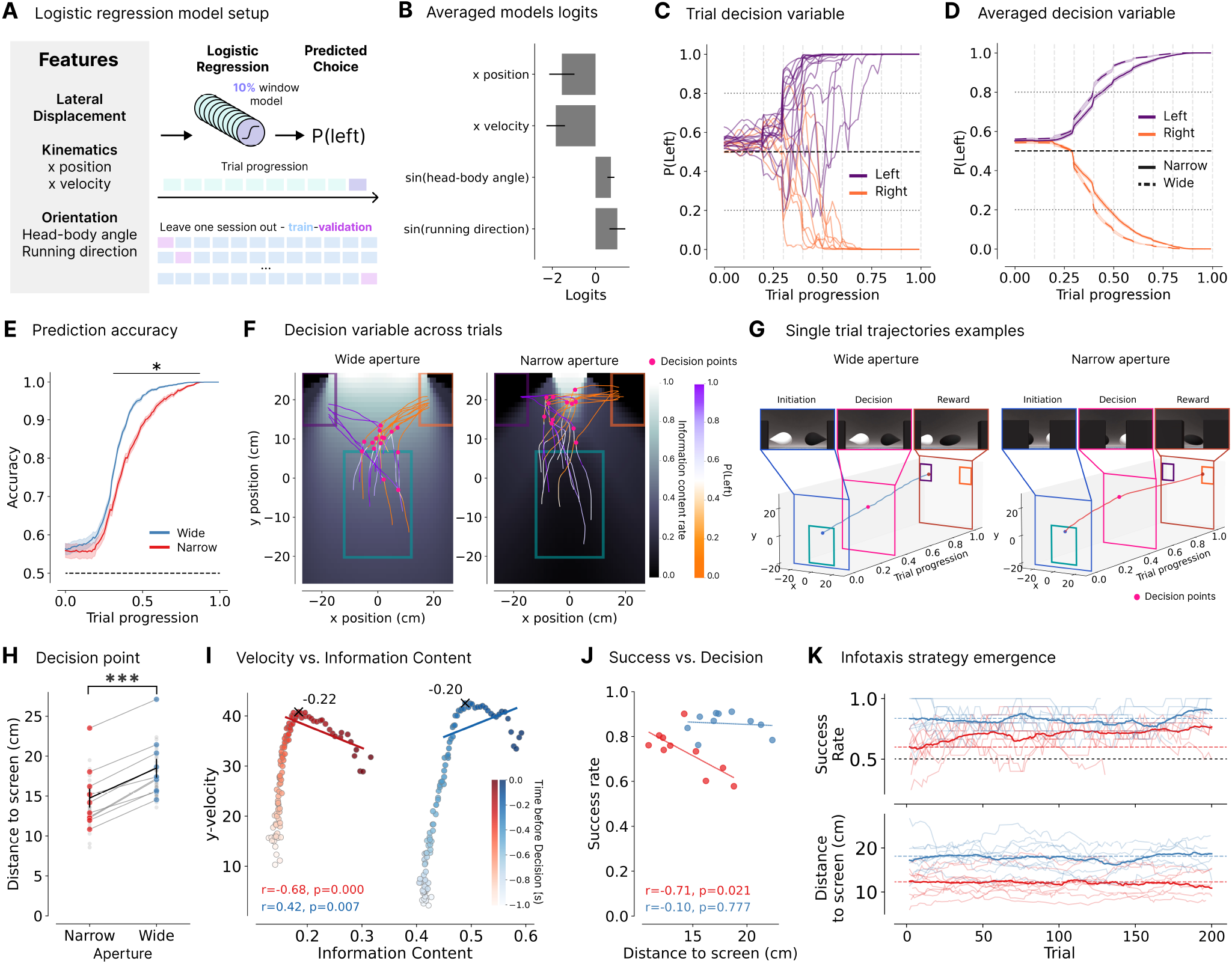
Behavioral features reliably define a dynamic choice variable revealing that, under occlusion, mice rapidly shift decision points to more informative viewpoints, correlating with better performance. **A**: Logistic regression setup. Four kinematic features (lateral position, velocity, head-body angle, and running direction) selected via BIC analysis to predict the choice side (Figure S4**A**). Ten independent models trained per session on non-overlapping 10% trial segments using leave-one-session-out cross-validation. **B**: Mean logistic regression weights (logits) *±* s.e.m. across sessions (n=43 sessions) and trial segments (n=10 segments; segment-specific weights: Figure S4**G**). Dynamic choice variable (*P* (*lef t*)) for **C**: example trials, and **D**: mean *±* s.e.m across trials and models per choice and aperture (n=43 sessions). Grey lines: decision threshold at a 20% confidence bound (STAR Methods). **E**: Model prediction accuracy per aperture. *∗* : FDR-corrected paired t-test with multiple comparisons across trial progression, significant aperture effect (*p <* 0.05) on 30-90% of trial progression, Figure S4**H**. **F**: Example trial trajectories colored by instantaneous dynamic choice variable; pink dots indicate decision points (20% confidence bound). Maps display object visibility across arena positions per aperture (STAR Methods). **G**: Single trajectory example, with stimuli at onset, decision, and reward. Decision at 36% (wide) and 44% (narrow) of trial. **H**: Decision point distance to screen (cm) per aperture and mouse. Means *±* s.e.m.: narrow: 13.79 *±* 0.44 cm, wide:17.54 *±* 0.42 cm (difference: 3.75 cm). *∗∗∗* : paired t-test, t(42) = 6.03, *p* = 3.56 *×* 10^*−*7^. Black: mean *±* s.e.m.; colored: per-mouse; grey: per-session. **I**: y-velocity vs. information content aligned to decision (from 1*s*) per aperture. Linear least-squares regression from *−* 0.4*s* to decision (*r*: Pearson correlation, *p*: Wald test p-value, *H*_0_: slope is zero): narrow: *r = −* 0.68, *p <* 0.001; wide: *r* = 0.42, *p* = 0.007. Peak velocity: *−* 0.22*s* (narrow) and *−* 0.20*s* (wide). **J**: Linear least-squares regression between success rate and decision distance to screen (cm) per aperture: narrow: *r* = *−* 0.709, *p* = 0.022; wide: *r* =*−* 0.103, *p* = 0.777. Per-mouse average across all sessions, regardless of the reward-drop. **K**: Success rate (above) and decision distance (cm) (below) across the first occluded session (200 trials, n=10 mice) using a 15-trial moving average. Dotted: first 15-trial baseline; solid: mean across mice; thin: individual mice. See also Figure S4.

This infotaxis behavior was correlated with higher success rates under the narrow-aperture condition but not under the wide-aperture one: mice that moved closer to the screen to make a decision on average were also the most successful on the occluded discrimination task, and this, even though all of them already performed well (above 0.7 success rate threshold; linear least-squares regression across all sessions, independent of the reward drop between conditions: narrow: Pearson’s *r* = − 0.709, Wald test, *p* = 0.022; wide: *r* = − 0.103, *p* = 0.777; Figure 4**J**; paired t-test between the *Closest* and *Furthest* mice quintile – 20% – groups per aperture condition: narrow: t(41) = −3.16, *p* = 0.0029, wide: t(41) =− 0.18, *p* = 0.86; Figure S4**K**). This suggests that moving the decision point closer to the screen is functionally relevant to task execution.

Interestingly, the immediate emergence of this strategy when mice were first exposed to occluders (in the very first dual occlusion test session; Figure 4**K**) suggests that they rapidly formed a robust internal model of the environment. Rather than gradually exploring or purely trial-and-error learning, they were able to apply this internal model efficiently, directly adjusting their trajectories and decision points to maximize information even under novel occlusion conditions. This indicates that mice can perform infotaxis not only re-actively, but also proactively, leveraging an internal representation of the task structure to guide efficient exploration and decision-making from the very first exposure.

### Infotaxis behavior correlates inversely with the amount of visible object area

The dual-occlusion task demonstrated that mice modulate their trajectories and decision location when visual information is limited. However, this binary manipulation could not reveal whether their information-seeking strategy scales continuously with sensory uncertainty. We therefore developed a second occlusion task with five uniformly presented occlusion levels, ranging from fully visible objects (99%) to fully hidden objects (0%) when the animal starts the trial from the center of the initiation area (Figure 5**A**; balanced aperture condition appearance, paired t-test *p >* 0.05, Figure S5**A**; Video S2). Across all three laboratories, mice performed the task with high accuracy. We observed a significant performance drop as the aperture narrowed, reflecting the gradual increase in task demand as visual uncertainty increases (FDR-corrected paired t-tests: 99% and 72% object visible conditions significantly different from the other conditions; Figure 5**B**, Figure S5**B**), yet they could modulate their behavior on a trial-by-trial basis to reliably report the location of the correct object without a population-level significant bias across all aperture conditions (all aperture condition choice bias indices remained close to zero – i.e., no bias, single-condition t-test vs. no bias: all *p >* 0.05; Figure S5**C**).

**Figure 5.**
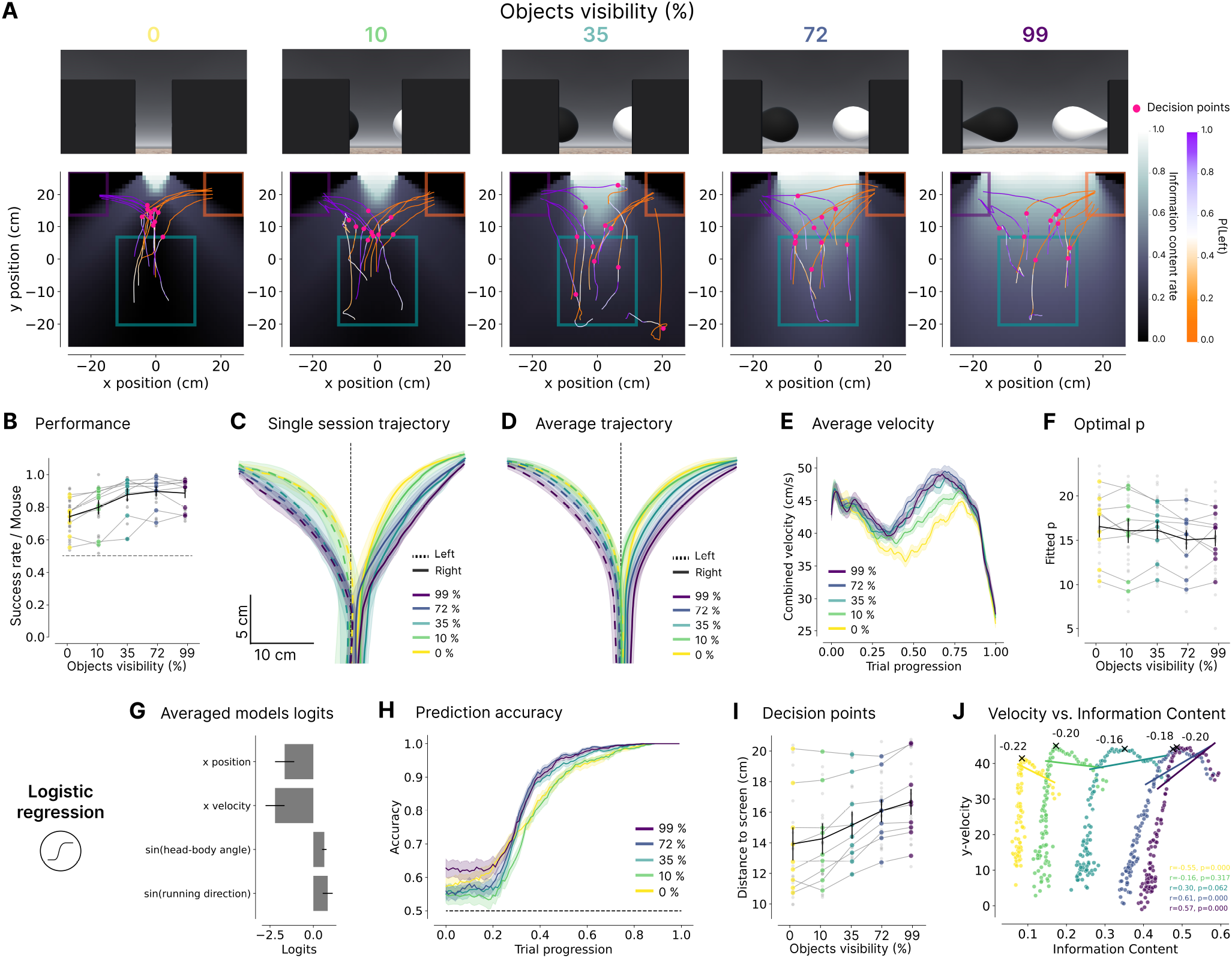
The amount of information available to the mouse correlates inversely with infotaxis behavior. **A**: Object visibility across five logarithmically distributed aperture levels in the multi-occlusion task, ranging from 99% to 0% of the objects visible (Above) and example trial trajectories colored by the instantaneous dynamic choice variable (pink: decision points). Information maps show object visibility across arena positions per aperture. **B**: Sucess rate per aperture condition and mouse. Repeated-measures ANOVA, significant aperture effect: F(4, 96) = 37.52, *p <* 0.0001. FDR-corrected paired t-test: *p <* 0.00025 for all but 10-99%, 0-35%, and 0-10% pairs; Figure S5**B**. Mean *±* trajectory s.e.m. (shaded) per aperture and choice for **C**: individual example mouse (Nightingale, s=4 sessions; Figure S5**D**) **D**: mean across mice (n=9 mice, s=25 sessions). FDR-corrected paired t-test with multiple comparisons across y bins (0-19 cm in the arena), significant aperture and choice effects (*p <* 0.05) between narrower and wider aperture conditions; Figure S5**E**. Midpoint to screen shown; dashed line: x-axis center. **E**: Mean velocity *±* s.e.m. per aperture across trial progression. FDR-corrected paired t-test with multiple comparisons across trial progression, significant aperture effect (*p <* 0.05) between narrower and wider aperture at 40-80% of the trial; Figure S5**F**. **F**: Optimal *p* parameter from single-trial L_*p*_ curve fits. Repeated-measures ANOVA, significant aperture effect: F(4, 96) = 3.14, *p* = 0.018. FDR-corrected paired t-tests: non-significant for all but the 0-72% pair; Figure S5**G**. **G**: Mean regression logits *±* s.e.m. across sessions (n=25 sessions) and trial segment (n=10 segments). **H**: Mean model prediction accuracy *±* s.e.m. per aperture. Two-way repeated-measures ANOVA, significant effect of aperture: F(4, 96) = 10.92, trial progression: F(49, 1176) = 428.47, and interaction: F(196, 4704) = 4.35; all *p* < 0.0001. **I**: Decision distance to screen (cm; 20% threshold) per aperture. Repeated-measures ANOVA, significant aperture effect: F(4, 96) = 39.61, *p <* 0.0001. FDR-corrected paired t-tests, significant differences for all but 0-10% and 72-99% pairs; Figure S5**R**. **J**: y-velocity vs. information content aligned to decision (from *−*1*s*) per aperture. Linear least-squares regression from *−*0.4*s* to decision (*r*: Pearson correlation; *p*: p-value), and peak velocity timing. For **B, F**, and **I**: Black: mean *±* s.e.m.; colored: per-mouse; grey: per-session. See also Figure S5 and Video S2.

To assess the behavioral correlates of information-seeking under varying occlusion levels, we analyzed the average trajectories per session (Figure 5**C**, Figure S5**D**). For all aperture widths, mice showed the same characteristic J-shaped trajectories as in the dual-occlusion task. The steepness of the curve varied systematically, forming a gradient from shallow (wider apertures) to steep (narrower apertures). Across all mice, we observed significant differences in the averaged trajectories between the narrower and wider aperture conditions (FDR-corrected paired t-test: *p <* 0.05 for pairs 0-99% and 0-72%, 28.3 cm away from the screen onward, and 0-35%, 10-99%, 10-72%, 10-35%, 16.2 cm away from the screen onward; Figure 5**D**, Figure S5**E**).

Consistent with increased deliberation, kinematic analyses revealed significantly lower velocities during the 40–80% portion of trials under narrow aperture conditions (FDR-corrected paired t-test: *p <* 0.05 for 0-99%, 10-99% and 35-99% pairs in 40-80% of the trial; Figure 5**E**, Figure S5**F**). Additionally, the optimal *p* parameter from fitting L_*p*_ curves (repeated-measures ANOVA: significant aperture effect: F(4, 96) = 3.14, *p* = 0.018, FDR-corrected paired t-tests: non-significant for all but the 0-72% pair; Figure 5**F**, Figure S5**G**) and trial tortuosity (repeated-measures ANOVA: significant aperture effect: F(4, 96) = 11.85, *p* < 0.0001, FDR-corrected paired t-test: *p <* 0.009 for all but 72-99%, 10-35%, 0-35%, and 0-10% pairs; Figure S5**H**) were both more elevated in narrower aperture trials, reflecting more circuitous paths.

The logistic regression framework trained to decode choice from lateral displacement behavioral features throughout trial progression (Figure 5**G**; Figure S5**I:P**) replicated the patterns observed in the dual-occlusion task. In particular, the model reached high confidence later in the trial for narrower apertures (two-way repeated-measures ANOVA, significant effect of aperture: F(4, 96) = 10.92, trial progression: F(49, 1176) = 428.47, and interaction: F(196, 4704) = 4.35; all *p* < 0.0001; Figure 5**H**; model accuracy: Figure S5**Q**), decision points were significantly closer to the screen under higher occlusion (repeated-measures ANOVA, significant aperture effect: F(4, 96) = 39.61, *p <* 0.0001; FDR-corrected paired t-test: *p <* 0.004 for all but 0-10% and 72-99% pairs; Figure 5**I**, Figure S5**R**) and instantaneous velocity was modulated by the corresponding available information (Figure 5**J**). Collectively, we observed that mice can perform active visual sensing and are behaviorally sensitive to relatively small changes in visual occlusion. Critically, we observed the same trends on the black target experiment (Success rates: 0%: 0.741 ± 0.019, 10%: 0.825 ± 0.030, 35%: 0.906 ± 0.021, 72%: 0.961 ± 0.016, 99%: 0.944 ± 0.016; Trajectories: two-way repeated-measures ANOVA, significant effects for aperture: F(4, 36) = 4.25, *p* = 0.0064, and spatial bin: F(24, 216) = 135.89, *p* < 0.0001, interaction not significant: F(96, 864) = 1.03, *p* = 0.4042, Combined velocity (m/s): two-way repeated-measures ANOVA, significant effect of aperture: F(4, 36) = 8.02, *p* = 0.0001, time bin: F(99, 891) = 36.20, and interaction: F(396, 3564) = 4.93; all *p* < 0.0001; Decision distance to screen: 0%: 13.785 ± 0.688, 10%: 14.765 ± 0.612, 35%: 16.407 ± 0.664, 72%: 17.583 ± 0.924, 99%: 17.385 ± 0.685; Figure S5**S**).

## Discussion

Here, we present evidence that mice engage in visual active sensing as defined by Yang et al. (13), who outlined three theoretical hallmarks: task-dependent movements, sensor- and actuator-dependent strategies, and history-dependent actions. Accordingly, mice robustly perform visual infotaxis. They move to acquire new views of the visual scene to overcome visual uncertainty and perform the task more accurately. Interestingly, this behavior is similar to the “peeking” observed in a predator avoidance task (45), where mice quickly move to acquire visual information while minimizing their own exposure. Furthermore, by analyzing the trajectories, we are able to quantitatively assess that both the decision point and the changes in behavior were dependent on the amount of visual information available, demonstrating sensor-dependent changes in behavioral strategies. The transition from more direct paths to more flexible ones as visual uncertainty increases – notably with the presence of “changes of mind” trials – aligns to active sensing strategies observed in other modalities, such as olfactory infotaxis, where animals switch from stereotyped tracking to sensory-driven search to resolve spatial ambiguity (46). Lastly, we show that these movements are not merely reactive to information accumulation but are learned over training to converge toward J-shaped trajectories and lead to more informative viewpoints. This directly increases task-relevant information and correlates with higher success rates. Notably, behavioral adaptations to sudden occlusions immediately upon their first presentation suggests that they can leverage a learned understanding of the task’s underlying structure to strategically sample information when faced with novel challenges.

Previous freely moving virtual reality platforms, such as FreemoVR (47) and related open-field systems (e.g. 48), demonstrated that unrestrained rodents can interact with perspective-correct environments and exhibit naturalistic behaviors. More recent immersive head-mounted approaches, including MouseGoggles (49) and Moculus (50), have further expanded the available visual field through stereo or panoramic displays though they typically constrain translational locomotion. The freely-moving virtual reality system developed here bridges these approaches and enables quantitative investigation of a range of visual processing in which fully manipulable input is dependent on an animal’s location or movement. This includes aspects of 3D perspective, motion parallax, and figure-ground segregation. By accurately coupling the visual scene to the animal’s pose and movements, the system reproduces realistic spatial and temporal dynamics of a physical environment, while maintaining experimental control of the rendered scene. It also provides a significant benefit over standard head-fixed virtual reality systems by allowing naturalistic head and eye movements – key aspects of active sensing, driving vestibular signals essential for the brain to incorporate information about self-movement (51).

A key next step is to resolve the neural dynamics that support this strategy, spanning the full sensorimotor loop from retina to motor output. Neural representations are shaped by the interaction between sensory input, motor commands, task structure (e.g., movement costs, reward contingencies), and intrinsic computational limits of neural circuits (52). Early visual areas are expected to reflect stimulus-driven signals, whereas downstream motor regions encode action plans. How these constraints are progressively integrated across the pathway to generate commitment and action, however, remains largely unknown – a systems-level question that our paradigm is uniquely positioned to address (53).

Normative frameworks such as Bounded Rationality propose that agents operate under representational constraints, limiting what information is retained and how it is encoded (54, 55). Different constraints should produce distinct signatures in both behavior and neural activity, including systematic patterns of choices and errors (56). By combining large-scale neural recordings with our task, it will be possible to link behavioral strategies to population coding principles, test competing models of representational limits, and determine how distributed circuits construct and transform latent variables that guide active sensing and decision-making.

## Resource Availability

### Lead contact

Requests for further information and resources should be directed to and will be fulfilled by the lead contact, Mackenzie Weygandt Mathis.

### Materials availability

This study did not generate new materials.

## Data and code availability

- The mouse behavioral data analyzed in this paper has been deposited at Zenodo (https://doi.org/10.5281/zenodo.19091270) and is publicly available as of the date of publication.
- All original code for this project has been deposited on GitHub (https://github.com/MMathisLab/FreelyMovingVR4Mice).
- Any additional information required to reanalyze the data reported in this paper is available from the lead contact upon request.

## Acknowledgments

We acknowledge funding from the NIH Brain Initiative U01 grant no. UF1-NS126566-02 (CN, AT, MWM, XP). CB and TS are co-first authors and may list their names in either order.

## Author Contributions

Conceptualization, M.W.M., A.T., C.N., and X.P.; Methodology, T.S., C.B., M.W.M., A.T., C.N., X.P., P.F., K.F, L.F., R.F., L.N., K.P., Z.T., and C.C.; Software, T.S., M.P., C.B., and L.B.; Simulation, C.B., and R.K.; Investigation, C.B., T.S., L.B., C.C., Y.L., K.J., R.F., L.N., K.P., and Z.T.; Writing-Original Draft, C.B., M.W.M., and T.S.; Writing-Editing, C.N., A.T., X.P., and K.F.; Funding Acquisition, C.N., M.W.M., X.P., and A.T..

## Methods

Key resources table.

| REAGENT or RESOURCE | SOURCE | IDENTIFIER |
| --- | --- | --- |
| **Experimental models: Organisms/strains** |  |  |
| Mouse: C57BL/6J: Wild-type (WT) | The Jackson Laboratory | RRID:IMSR_JAX:000664 |
| **Deposited data** |  |  |
| Mouse Behavioral Data | This paper | https://doi.org/10.5281/zenodo.19091270 |
| **Software and algorithms** |  |  |
| Python Programming Language v3.9—3.10 | Python Software Foundation (57) | RRID: SCR_008394; https://www.python.org/ |
| DataJoint | Yatsenko et al. (58) | RRID:SCR_014543; https://www.datajoint.com/ |
| DeepLabCut v2.2+ | Mathis et al. (59), Nath et al. (60) | https://www.mackenziemathislab.org/deeplabcut |
| *DeepLabCut-Live!* v1.0.4 | Kane et al. (41) | https://github.com/DeepLabCut/DeepLabCut-live |
| SuperAnimal-TopViewMouse | Ye et al. (61) | https://huggingface.co/mwmathis/DeepLabCutModelZoo-SuperAnimal-TopViewMouse |
| FreelyMovingVR4Mice | This paper | https://github.com/MMathisLab/FreelyMovingVR4Mice |

### Experimental Model and Study Participant details

Behavioral data was collected from 10 C57BL/6J mice (wild-type) across three laboratories, respectively 6 in the MLAI (M-Lab of Adaptive Intelligence), 2 in the Tolias laboratory, and 2 at the Niell laboratory. All mice underwent the same behavioral protocol, including training, and the two test phases (dual and multi-occusion tasks). They were housed on a reversed 12-hour light-dark cycle from 9 am to 9 pm. All experiments took part within the dark phase of that cycle. Both male and female mice were used and were between the ages of 8 to 20 weeks (MLAI: 5 females, 1 male; Tolias laboratory: 2 males; Niell laboratory: 2 males). Mice were placed on a controlled water access protocol for training, where they were maintained above 90% of their original body weight and were given no less than 1.5 mL of water per day. These experiments were carried out using the same protocol in three laboratories: the Tolias laboratory, Baylor College of Medicine (Texas, USA) and Stanford University (CA, USA) under the BCM OLAW Assurance number D16-00475 and Stanford Animal Welfare Assurance number D16-00134, the MLAI, Campus Biotech-EPFL (Geneva, Switzerland) according to the Geneva canton license number GE367, and the Niell laboratory, University of Oregon (Oregon, USA) under the protocol number AUP-21-21.

### Method Details

#### Behavioral experiments

The same experimental protocol was followed in all three laboratories that carried out the experiments (Figure S2**A**).

#### Habituation and behavioral training

Prior to habituation and training, WT mice were placed on a controlled water schedule in their home cage. They were then habituated to the experimenter for a period of 3 to 5 consecutive days. On the first day, the experimenter placed their hand in the cage for 15 to 20 minutes to allow the mouse to acclimate to the experimenter’s scent. On the second day, the experimenter gently picked up the mouse using a plastic tube and then returned it to the cage. This was repeated multiple times during 15-20 minute sessions. On the third day, the mice were moved from the plastic tube onto the experimenter’s hands multiple times, allowing them to freely explore the hands. If a mouse showed signs of stress at any stage, the day was repeated until the mouse was comfortable. Following this habituation period, mice were habituated to the virtual reality arena by allowing them to freely explore the box for 30 minutes. To encourage mice to associate lick ports with rewards, a small water reward of 5 *µ*L was given whenever mice were in close proximity to the right or left lick ports. This water reward was manually triggered by the experimenter using the manual water task on the `vr4mice` GUI.

The virtual world is rendered in the real-world arena on a 520-mm-wide screen. Once habituated, mice were trained in three phases: *1*.*1) detection without a velocity threshold, 1*.*2) detection with a velocity threshold*, and *2) discrimination*. Mice progressed through stages upon meeting a predefined criterion, which required completion of more than 125 trials in 60 minutes with a success rate greater than 70%. During the detection phase (*1*.), a single object, a white teardrop, appeared randomly on the right or left side of the virtual world with a probability of 0.5. The mouse reported the location of the object by moving to the lick port on the side of the object. In the first phase of the detection task (*1*.*1*), we did not enforce a strict trial initiation threshold: mice had to stay in the trial initiation area for at least 100 ms with a velocity lower than 20 Unity units per second (~ 67.5 cm/s) to start the trial. Then in the second phase of the detection (*1*.*2*), the velocity threshold was reduced to 10 Unity units per second (~ 33.75 cm/s), and the time limit increased to 250 ms. In the discrimination phase (*2*.), a distractor object, a black teardrop, was presented on the opposite side of the white teardrop, with still a probability of 0.5 to appear on one side or the other and a trial initiation threshold similar to *1*.*2*.

#### Discrimination with occlusion

In the discrimination with occlusion test, occluding walls were placed in front of the objects of interest. These occluders consisted of two walls separated by a central aperture, allowing the mice to observe the objects through the gap. The extent of occlusion varied with the aperture width, causing the objects of interest to be either partially, fully visible, or fully occluded, varying the amount of information the mice could gather about the objects from a stationary position.

In the dual-occlusion test, mice observed objects through a wide or narrow aperture (Figure 3**A**), restricting the visible portion of objects of interest. When standing at the center of the trial initiation area, 99% of the objects was visible for the wide aperture condition (5.6 cm walls on each side, with an aperture of 40 cm), compared to 1% for the narrow one (19 cm walls on each side, with an aperture of 15 cm). The aperture condition was randomly selected for each trial using a uniform distribution.

After completing this task for five sessions, the mice were then tested on the multi-occlusion task, with five different aperture conditions ranging from 0% to 99% objects visibility on a logarithmic scale (0%, 10%, 35%, 72%, and 99% of the objects visible; Figure 5**A**), also from the center of the trial initiation area. This second task was also conducted for five sessions.

#### Inclusion criteria

To ensure that only sessions with sufficient task engagement were analyzed, we applied a post hoc inclusion criterion to test-day sessions (dual or multi-occlusion). Mice had to have completed more than 125 trials with a success rate greater than 70%, and their reward rate could not drop by more than 25% between the least and most occluded conditions for the session to be included. In total, across ten mice, we ended up with 43 sessions for the dual-occlusion test and 25 sessions on the multi-occlusion test (with one mouse that performed no session above the criterion). For the analysis correlating decision-point distance with success rate, all sessions with more than 125 trials and a total success rate greater than 70% were included regardless of the reward-drop criterion. This approach was taken to capture the full range of behavioral strategies across both high- and low-performing sessions.

### The virtual reality system

#### The arena

The virtual reality arena consisted of a transparent box measuring 52 × 52 cm. The arena was constructed using a base plate and four side panels, all made of PMMA (polymethyl methacrylate; Angst and Pfister). The base plate measured 520 × 520 × 5 mm, while each side panel measured 525 × 360 × 5 mm. These side panels were securely attached to the base using aquarium primer and glue (Coltogum). Inside the box, an additional floor plate with an anti-glare adhesive coating (Gardinia) was placed to minimize reflections within the arena. Sandwiched between the inner floor plate and the outer floor plate was a sheet of light diffuser (LEE filters), which allowed for even illumination of the arena. This inner floor plate could be easily removed for cleaning between sessions.

#### The mounting cage

The plastic box was mounted onto a cubic metal frame constructed from 25 mm aluminum optical rails (Thorlabs). The frame was built using eight 540 mm rails for the top and bottom faces of the cube and four 550 mm rails for the vertical sides. These rails were connected at the corners using Thorlabs corner cubes, with additional right-angled brackets added for extra stability. The transparent box was attached to this frame using custom 3D-printed top and bottom right-angled brackets. These brackets were secured to the optical rails with drop-in T-nuts and 1/4”-20 low-profile screws. The 3D-printed brackets also supported three computer monitors (SB241Y, Acer, 540 mm × 316.5 mm) coated with 0.9 neutral density filters (LEE filters) to achieve a maximum luminance of 10 cd/m^2^.

#### Camera and computer

Above the cage, a high-speed USB camera (DMK-37BUX28, Imaging Source) was mounted using a right-angled bracket attached to a cross-beam assembly consisting of one 540 mm beam and two 200 mm beams. The camera, equipped with a 3.5 mm EFL, F/1.4, 1/2” lens (Navitar), was positioned centrally over the arena. The cross-beam could be adjusted vertically, allowing the camera to have a complete field of view of the arena floor. The camera was connected to a Lambda vector Windows 10 workstation equipped with an Intel Core i9-10900X processor, 2 × RTX 3080 GPUs, 128 GB RAM, and two hard drives (1 TB for the operating system and 2 TB for data storage). This system managed camera capture, pose estimation, and generated the signals for water delivery.

#### Experimental environment acoustic stability

To ensure behavioral performance was not influenced by dynamical acoustic cues (e.g., electronic noise such as “coil whine” from the computer monitors due to changing brightness for instance), ambient sound was recorded during a representative 10-minute session using an iPhone 12 (Micro-Electro-Mechanical Systems microphone array; 44.1 kHz sampling rate) and analyzed post-hoc for spectral and temporal stationarity using the Librosa and SciPy libraries. The acoustic environment was characterized by a stable, low-frequency distribution. No periodic pulses, vertical spikes, or “blocks” of noise correlating with task-related screen updates was visible. The sharp horizontal cutoff at 16kHz represented the frequency response limit of the recording device (Figure S1**H** – Left-Top). The Root Mean Square (RMS) of sound pressure levels was extracted to monitor the noise for sustained step-functions or repeated task-locked pulses (Figure S1**H** – Left-Down). It revealed a highly consistent baseline noise floor. High-amplitude spikes correspond to mechanical artifacts, such as mouse or experimenter’s movement, rather than task-locked stimulus onsets. The absence of sustained amplitude shifts indicates that the computer hardware did not produce volume-based cues during task intervals. Power Spectral Density (PSD) was calculated for discrete 2.5-minute segments distributed across the session using Welch’s method to identify potential frequency shifts (Figure S1**H** – Right). The absence of unique narrow-band spectral peaks in any single segment confirms that no new dynamical frequencies (e.g., variable coil whine) were introduced during the execution of the behavioral task.

#### Water delivery and lick ports

Two lick ports were placed on the left and right sides of the arena, each located 5 cm behind the front screen. These lick ports consisted of custom-made 3D-printed boxes, each with a hole near the top to allow a length of water tubing to pass through. Inside each lick port, the water tube was securely fastened to the top of a steel pedestal (Thorlabs) using a separate 3D-printed attachment. This design ensured that the mice could not pull the tubing through the lick port and added enough weight to prevent the lick port from being knocked over by the mouse.

The water delivery system was controlled by a Teensy 4.0 microcontroller via a TTL (Transistor-Transistor-Logic) signal that was sent to the valve, with the duration of the TLL specifying the time in ms for the valve to stay open. Upon receiving a signal from the computer, the microcontroller would open one of the two solenoid valves, corresponding to either the right or left lick port. When a valve was opened, water would flow from a reservoir syringe (20 mL) through the tubing to the lick port, providing the mouse with a water reward of 3-5 *µ*L.

To ensure that the valve’s open time matched the desired droplet volume, a calibration curve was created for the lick ports. The valve’s open time was adjusted in 100 ms increments, ranging from 100 ms to 1000 ms. At each interval, the droplet size was measured either by collecting the droplet in a weigh boat and weighing it on a precision scale or by using a pipette. These measurements were used to plot a calibration curve, which identified the time range that produced a droplet volume of 3-5 *µ*L.

Prior to each experimental session the water systems tubing was cleared of any air bubbles by inserting a syringe plunger into the syringe reservoirs. Air bubbles were then pushed through the system by giving 5 × 1000 ms duration pulses, using the manual water task in the vr4mice GUI. Following this the plungers were removed and the syringes were topped up to 20 mL, ensuring the same water pressure between sessions.

### Software

#### *DeepLabCut-Live!* pose estimation and online kinematics

To provide closed-loop visual feedback to the mouse, its head position within the arena was estimated using the DeepLabCut-Live GUI (41), and used to render the corresponding viewpoint in Unity. DeepLabCut-Live utilizes a camera processor that receives images of the arena from the top camera at 100 Hz; complete parameters are listed under *Camera and DeepLabCut-Live parameters used*. The pose of the animal is inferred using the pre-trained TensorFlow SuperAnimal-TopViewMouse model (61) (Figure S1**A**), video adapted on a top-view video of our arena. This model, which has been optimized across multiple freely moving mouse datasets, yields 27 keypoints over the body of the mouse when viewed from above. From these keypoints, a custom-built *DeepLabCut-Live!* processor Python script computed kinematic features including the x- and y-position of the head, the running direction and the head angle relative to the body (head-body angle) (Figure S1**B-C**). The x- and y-position of the head were calculated by taking a weighted average of the eight keypoints corresponding to the head, namely `head_midpoint, right_ear_tip, left_ear_tip, right_ear_base, left_ear_base, right_eye, left_eye`, and `nose`:

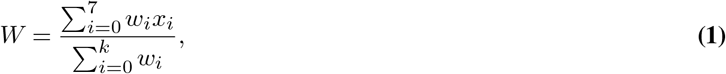

where *w*_*i*_ is the confidence score of keypoint *i* (i.e., DeepLabCut likelihood) and *x*_*i*_ is the x-axis position of keypoint *i* (same for y-axis). If the mean confidence of all head keypoints was below 0.6, the previous high-confidence position was used instead. This prevents low-confidence estimates from affecting head position in frames where the head is occluded, such as when grooming, or proximity to arena walls.

To compute the mouse running direction, we defined a body axis vector from the tail base keypoint to the neck keypoint. The running direction was the angle (in degree) between this body axis and the horizontal axis (x-axis). We then adjusted this angle so that 0° aligned with the main monitor, with values ranging from − 180° to 180°. To obtain the head-body angle, we first calculated the head axis as the vector from the neck keypoint to the nose keypoint. The angle between the head and body axes then gave the head-body angle (Figure S1**B**). These kinematic features were sent to the `vr4mice` Python framework via a UDP (User Datagram Protocol) socket at a rate of around 70 Hz to determine when to trigger the trials.

Camera and DeepLabCut-Live parameters used.

| Camera processor |  |
| --- | --- |
| camera_name | Imaging_source |
| resolution | (720, 540) |
| exposure | 0.01 |
| rotate | 0 |
| crop | (20, 530, 90, 620) |
| fps | 100 |
| display | True |
| display_resize | 0.7 |
| DeepLabCut-Live inference |  |
| model_name | superanimal_topview |
| model_type | base |
| cropping | None |
| dynamic_cropping | False, 0.5, 10 |
| resize | 1.0 |
| mode | optimise_latency |

#### VR4Mice Python framework

To run the virtual reality system, we created a custom Python framework called `vr4mice`. This framework acts as an interface between the incoming DeepLabCut-Live data, and Unity and controlled the valves for water delivery via a Teensy 4.0. In addition, the `vr4mice` software allows the user to edit task parameters (e.g., trial structure or task type) and save the data through a GUI.

`vr4mice` continually reads data from the DeepLabCut-Live GUI via the UDP socket on a separate thread and adds the incoming data to a buffer. The main thread periodically checks the buffer for the latest entry and reads it whenever Unity is ready to process a new frame. The data is then passed through a “one-euro” filter, a fast first-order low-pass filter with a cutoff that adapts to the sampling frequency, to reduce jitter in the signal (62). It is then converted from pixel space (720× 540 pixels) to game space (18 × 8 game units) using the numpy `interp` function (63) and sent to the Unity game. The position in the arena, inferred from the DeepLabCut keypoints, is used to render the corresponding viewpoint and the running direction and head-body angle are used to initiate the trials in the game. At each iteration, the state of the mouse in the virtual game, which consists in its position and the reward status, is returned. If a reward was given in the Unity game, then `vr4mice` sends a signal via a serial port to a Teensy 4.0 to control the corresponding lick port.

#### The Unity game

The Unity game environment was constructed on a plane measuring 18 × 8 game units, which served as the floor of the virtual arena. The position of the real mouse’s head was transformed into the Unity coordinates and used to control a Unity empty object, representing the virtual mouse within the game, through the Unity Machine Learning Agents Toolkit, the ML-Agents package (64). A camera object was attached to this empty object, and its frustum was aligned with a projection plane (see the *Off-axis projection* Methods below for detail). This setup allowed objects of interest, designed with the software tool Blender (65) and randomly spawned behind the projection plane, to be viewed from the mouse’s perspective as it navigated the arena displayed on the main monitor.

We define a trial initiation area in the arena as a 24 × 18 cm (8× 4 Unity units) zone centered on the x-axis. It is positioned 7.75 cm from the back wall and 21.25 cm from the screen. A trial initiates only when the mouse remains below a velocity threshold of 10 Unity units/s for at least 250 ms (20 Unity units/s for at least 100 ms at the start of training). As the game arena plane is not anisotropic, this corresponds to a velocity threshold of 33.75 cm/s on the y-axis and 15 cm/s on the x-axis, respectively 67.5 cm/s and 30 cm/s, considering that the mice direct themselves mostly on the y-axis at trial initiation. Additionally, the mouse’s head angle needed to be oriented toward the main monitor within a ± 90^°^ angle to the perpendicular axis. This threshold, which does not account for the precise angle of the screen from the mouse position, was sufficient given the central placement of the start box. If those criteria were met, two objects of interest were spawned on the left or right of the projection plane on the screen. The mouse could then report the location of an object of interest by going to one of the two lick ports. To achieve this, two report boxes were implemented in the Unity game, with their positions corresponding to the lick ports’ location in the physical arena. If the virtual mouse stayed within the correct report box for at least 100 ms, a reward signal was sent to the `vr4mice` software via a socket. If the mouse reported incorrectly, no reward signal was sent. After the mouse had reported, the Objects of Interest disappeared from the projection plane and the next trial could be triggered by re-entering the trial initiation box.

#### Off-axis projection

In typical Virtual Reality systems, the subject’s visual field is centered within the projection screen and the virtual environment is rendered using an on-axis projection, where the camera frustum is symmetric on all sides relative to the center. The camera can be moved and rotated to follow the subject’s line of sight, maintaining a consistent and immersive experience. In our proposed system, the subject moves freely, changing the viewing angle relative to the screen. Consequently, an on-axis camera would fail to replicate certain aspects of natural vision, such as accurate motion parallax, leading to a less immersive experience. To address this, Cave Automatic Virtual Environment Virtual Reality (CAVE-VR) systems typically employ an off-axis projection camera instead (66). This approach adjusts the projection matrix to match the subject’s specific viewpoint, ensuring that virtual content is accurately rendered in alignment with the user’s perspective. It also maintains correct motion parallax, where distant objects appear to move less in response to head movements than closer objects, enhancing the realism of the rendering and making objects appear to be 3-dimensional.

To implement the off-axis projection, a projection plane was added to the Unity game. This projection plane acted as the near-clipping plane for the camera, meaning that any objects between the mouse and the projection plane would not be rendered. The central point of the camera perspective was fixed in the center of this projection plane. The corners of the frustum were fixed to the corners of the projection plane. The camera could then be moved around in Unity game coordinates and its view point rendered from the mouse’s perspective (Figure 1**B**; Figure S1**D**).

#### Latency testing

To measure the latency of the virtual reality system, the latency of the full system was measured along with the latencies for the DeepLabCut-Live and Unity steps (Figure S1**E:G**). A binary signal was generated in the DeepLabCut-Live processor so that it generates a binary signal *S*:

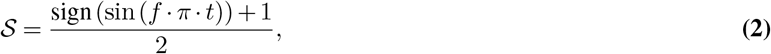

where *f* is the pulse frequency and *t*, the current time. Setting *f* to 5 Hz, the pulse of the binary signal *S* flips every 200 ms. This signal *S* was saved alongside the camera timestamp and the timestamp at which DeepLabCut-Live had completed frame processing. *S* was initiated 10 seconds after the Unity game started, ensuring that it had time to load before the signal started. It was then sent to the Unity game via a UDP socket and rendered as a flashing black-and-white square in the top right corner of one of the side screens. A photodiode attached to this screen recorded the rendered signal *R* via a Teensy 4.0. This signal *R* was sent back to the computer via USB to be read on a separate thread, time-stamped within the DeepLabCut-Live processor script at 1000 Hz and saved alongside the data for the originally generated signal *S*.

To measure end-to-end latency (Figure S1**F**), the photodiode signal was *R* first low-pass filtered at 60 Hz to remove high-frequency noise. This filtered signal was then normalized between 0 and 1 and binarized using a 0.5 threshold. Rising edge timestamps from this binarized signal *B* were compared to those of the generated *S* signal and their difference corresponds to the system’s end-to-end latency (Figure S1**G**). To determine the latency introduced by DeepLabCut-Live – namely frame capture, pose estimation, and keypoint averaging – we measured the time difference between the timestamp recorded at frame capture and the timestamp after computing kinematic features. The latency introduced by Unity was then calculated by subtracting the DeepLabCut-Live latency from the end-to-end latency (Figure S1**E**).

### Behavioral analysis

#### Quantifying information content across the arena

The information available to the mouse depends on its position in the arena, and how much of the two target objects were visible from that position. Here we defined information available as the objective visual evidence available in the environment (the surface of both objects), without making assumptions about the mice’s specific internal heuristic to solve the task (i.e., detection of the target object, or inference from the distractor object). For simplicity, we assumed these targets were circular. Each discrete point in the arena is treated as a visual sensor situated at a fixed elevation. The position of the target and aperture define the set of rays from the sensor that reach either target without intersecting the occluding walls. To calculate the unoccluded area of the targets visible from a given position, we compute the azimuthal angles subtended by unobstructed rays from each lattice point to both targets. These angular spans are converted to the corresponding circular-segment areas via 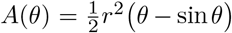. The scalar information available at each position (*x, y*) was then related to the visible areas as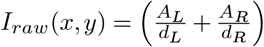, where *A*_*L*_ and *A*_*R*_ denote the visible segment areas of the left and right targets respectively and *d*_*L*_ and *d*_*R*_ are their corresponding Euclidean distances to the sensor. To faciliate comparison across conditions, the resulting information maps were min-max normalized to the range [0, 1] across all conditions: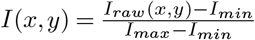, where *I*_*min*_ and *I*_*max*_ are the minimum and maximum values of *I*_*raw*_ across all information maps.

#### Off-line computation of kinematic and trial features

To analyze the DeepLabCut-Live data offline, keypoint traces were preprocessed to obtain traces that were free of keypoint jumps. This was achieved by removing all keypoints with a confidence value of less than 0.6. These low-likelihood keypoints were then inferred using neighboring keypoints using a linear interpolation. To reduce high-frequency jitter in keypoint tracking, the interpolated traces were smoothed using a Savitzky-Golay filter (SciPy (67)). To synchronize these traces with the Unity game data, the game data were first resampled to 50 Hz to correct for timing inconsistencies between frames. The DeepLabCut keypoint traces were then aligned by matching each timepoint to the closest one in the resampled Unity game recording. Considering that the Unity arena of dimensions 18 × 8 Unity units corresponds to a physical arena of 52 × 52 cm, and that screens are positioned 1 cm away from each wall, we interpolated the position of the animal with respect to the screens from the Unity coordinates to a physical space of 54 × 54 cm. We derived kinematic features such as the mouse’s head position, running direction and head angle relative to the body axis (head-body angle) similarly to what was implemented online for closed-loop feedback (Figure S1**B**). Running direction provides us with information on the spatial orientation of the mouse across the trial progression. Head-body angle serves as a measure of how attentional focus aligns with body movement, as supported by the activity of head-direction cells that are encoding the animal’s heading independent of body orientation (68). We also report the cosine and sine values for each angle to be used in the regression models, avoiding discontinuities and preserving directional structure. Cosine values are related to the forward/backward direction while sine values are related to the left/right direction in the arena. The x- and y-velocity in cm/s are computed from deriving the x- and y-position and used to obtain the combined velocity in cm/s. The initial position distribution heatmaps represent the spatial probability density of the animals’ starting coordinates, binned into a 3×3 grid. Each bin represents the mean occupancy for sessions and mice. We use this as an indicator of how consistent the mice were when approaching the start zone and how similar the initial stimuli were from trial to trial. We also tested for potential choice-predictive spatial biases by comparing the frequency of starts between Left and Right choices.

We also compute some trial metrics. To investigate whether previous trial outcomes influenced current choice behavior, we analyzed the side bias and decision stickiness. The side bias index detects persistent preferences for the left or right reward ports, while the decision stickiness index measures a potential tendency to repeat or alternate choices regardless of the outcome. Trial tortuosity is the trial path length (sum of the distances between all consecutive points) divided by the direct path from start to end position of the trial (distance between start and end points). From this, we defined change-of-mind trials as the 90^th^ percentile most tortuous forward trials. We classified a trial as “forward” if the minimum y-position was the initial position, hence excluding trajectories where the animal went toward the back of the arena during the trial. J-shaped trial rate is the rate of characteristic J-shaped trials, compared to wandering trials. We only include J-shaped trials, selected based on a duration (≤ 5 sec) and tortuosity (≤ 5) threshold, in our analysis, as an indication of the mouse’s engagement in the task. Time to report is the duration of the trial without the inter-trial interval, from trial initiation to the mouse getting to one reward port.

#### Training phase analysis

As mice required a variable number of days to complete training (Figure 2**C**), we assessed behavioral changes by selecting three representative sessions for each training phase (detection and discrimination). In the detection phase, the first and last of the detection without the velocity threshold and the last session of the detection with the velocity threshold. For the discrimination stage, the first, the penultimate, and the final session. These timepoints were chosen to approximate equivalent learning stages across mice. In the discrimination phase, some mice completed the phase in only two sessions, in which case no middle session was available. To account for this, statistical analyses were performed using a one-way ANOVA followed by Tukey’s HSD post-hoc test, excluding the middle timepoint when not present.

#### Trajectory analysis

For each session, we first preprocessed the data by *1)* removing the inter-trial interval, and *2)* removing the wandering trials to only keep the J-shaped trials. To characterize trajectories across occlusion conditions, the x- and y-position from individual J-shaped trials were first interpolated such that each trajectory comprised an equal number of time points (100 timepoints; trial progression). For each session, mean trajectories were computed by averaging the x and y coordinates across trials at each interpolated time point, separately for each aperture condition and choice direction (left or right). These session-level trajectories were then aggregated across mice to generate mean trajectories for each aperture. A similar procedure was used to calculate the other kinematics showed as a function of the trial progression.

For statistical comparisons, trajectories, running direction and head-body angles from right-choice trials were transformed by reflecting their x coordinates or angle across the midline (i.e., *x* → −*x*), aligning them with the left-choice reference frame.

#### Trajectories approximation with curves of constant L_p_ norm

To approximate the trial trajectories, each interpolated trial was fitted with a curve of constant L_*p*_ norm, which is a special case of the superellipse (Lamé curve) with equal horizontal and vertical scaling:

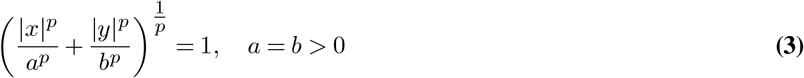

where *p* = 0 is the shape exponent that controls the curvature of the curve and *a* and *b* are scaling factors set to the maximum Euclidean distance per trial. Higher values of *p* produce more angular paths, axes-aligned paths that extend forward in the arena before turning toward the water ports, while lower values yield smoother, more direct trajectories, where mice move more directly to the water ports. For each trajectory, the optimal *p* parameter was estimated by fitting the L_*p*_ curve to the trials’ x and y position. So that they are comparable across trials, trajectories were first normalized by their maximum Euclidean norm:

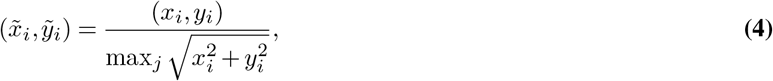

where *i* indexes points in the trajectory and *j* spans all timepoints. The L_*p*_ curve was the fitted by minimizing the discrepancy between the observed and predicted y-coordinates. Given a value of *p*, predicted y-values were computed from the normalized x-coordinates as:

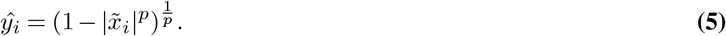

The optimal *p* was obtained by minimizing the sum of squared residuals:

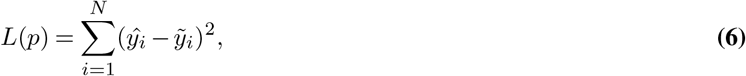

where *N* is the number of trajectory points. The resulting optimal *p* values were averaged across trials to quantify trajectory shape as a function of occlusion condition.

#### Black target contrast experiment

To ensure that the strategy employed to perform the task is goal-directed rather than reflexive attraction to the high-contrast white teardrop or low-contrast black teardrop compared to the gray background, we performed the inverse contrast scheme experiment, i.e., the target object is the black teardrop and the distractor is the white one. All other parameters were kept the same. We trained three female mice, in the MLAI. It took an average of 10.3 ± 0.3 days and a total of 1489 ± 106.81 trials to train them. We tested each mouse on 6 days of the dual-occlusion task, and 6 days on the multi-occlusion task. 6, 5, and 4 sessions respectively were found above the inclusion criteria, for a total of 15 sessions for the dual-occlusion task (Figure S3**Q**). 4, 3 and 3 sessions respectively were found above the inclusion criteria, for a total of 10 sessions for the multi-occlusion task (Figure S5**S**).

### Quantification and Statistical Analysis

#### Statistical Analysis

For trajectory and velocity analysis, a repeated measures two-way analysis of variance (ANOVA) was performed with aperture and y-bin location or trial progression as within-subject factors and session as the subject-level random effect (69). For the multi-occlusion task, variables for each aperture were compared using a repeated measures one-way ANOVA. Post hoc paired t-tests were conducted to further evaluate pairwise differences between aperture conditions. All resulting p-values were corrected for multiple comparisons using the Benjamini–Hochberg false discovery rate (FDR) procedure (70).

### Dynamic Choice Variable

#### Logistic regression

We implemented a time-resolved multi-model framework to infer mouse choice from behavioral features using logistic regression (scikit-learn; (71)). Rather than fitting a single global model, we partitioned each trial into ten non-overlapping temporal segments, each representing 10% of the interpolated trial, and trained an independent model for each segment. This approach accounts for the non-stationarity of behavioral features by allowing regression weights to adapt to the specific dynamics of each trial phase.

For each of the ten models, we used a leave-one-session-out (LOSO) cross-validation approach, training one model per session on the data from N-1 sessions and evaluated on the single held-out session. Model performance was reported as the mean per-timepoint prediction accuracy across trials and sessions for the given time segment. To quantify the decision process, we also reported the model’s probability estimates for left versus right choices as a function of trial progression, which we considered as a dynamic decision variable. These estimates correspond to the predicted probability of a left choice (*P* (left) = 1 indicating full confidence in a left choice; *P* (left) = 0 for a right choice) at each timepoint. These probabilities were computed on the held-out validation session of each model and subsequently averaged across trials and sessions.

#### Bayesian Information Criterion analysis

To identify the feature set that best captures the animals’ underlying decision dynamics, we evaluated the Bayesian Information Criterion (BIC) (72, 73) for ten regressor sets, corresponding to distinct behavioral hypotheses. First, the *Priors* set, composed of the initial positions (x and y) and the trial history, tested whether the choice was pre-determined by starting conditions and past outcomes rather than current trial dynamics. Then a *Kinematics* set, comprising position (x and y) velocity (x and y), assessed the extent to which the decision was explainable from basic movement parameters. To further isolate the specific drivers of this movement, we tested two reduced hypotheses: *Position only* and *Velocity only*. These sets evaluated whether the decision variable was primarily reflected in the animal’s spatial coordinates or its instantaneous movement vectors, respectively. The *Orientation* set included the running direction and head-body axis (sine and cosine) to evaluate if the decision was represented in a purely egocentric or directional frame rather than a spatial one. The

*Lateral Displacement* set solely comprised the features related to lateral movements (toward reward ports), namely x-position, x-velocity, and sine of head angle and running direction. The *Task-relevant* set expanded upon these by integrating kinematics and orientation with dynamic task features (trajectory tortuosity and length, and whether the trial was rewarded) to determine if the decision variable represented a synthesis of behavioral state and task context. Finally, the *Full State* set combined previously mentioned features, namely position, velocity, running direction, head-body angle, the dynamic task features and the prior features (initial position and trial history) to check whether pre-trial priors and online dynamics provide non-redundant contributions to the decision variable or if a more parsimonious model of current dynamics is sufficient to explain the behavior. The BIC is defined as

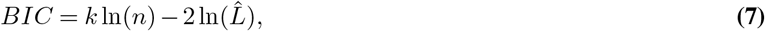

with *k*, the number of features used in the model, *n*, the number of observations (sample size) and 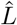, the maximized likelihood. For our binary choice prediction, the log-likelihood is calculated as ln 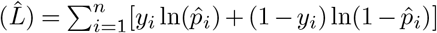 This metric rewards models that assign high probability to the animal’s actual choices while penalizing excessive parameterization; consequently, lower BIC values indicate a more parsimonious fit.

However, rather than seeking a model based only on aggregated goodness of fit across timepoints and trials, we identified the feature set that most effectively reflects the dynamic evolution of the decision process as the trial progress. Hence, we evaluated three distinct criteria: mean accuracy, mean BIC score, and the Coefficient of Variation (CV) of the BIC within the final 10% of the trial (*CV*_*end*_). The later serves as a metric for terminal stability, defined as:

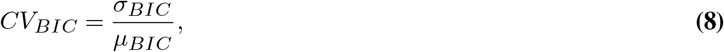

where *σ*_*BIC*_ and *µ*_*BIC*_ are the standard deviation and lean of the *BIC* values calculated across individual trials, respectively. To integrate these metrics into a single selection criterion, we perfromed a min-max normalization for each component to a scale of [0, 1]. We define a model efficiency score *S*, as the unweighted average of these normalized components:

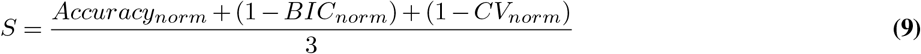

This score identifies the feature set that maximizes predictive power (high accuracy) and parsimony (low BIC) while ensuring the model reaches a stable, low-variance state as the animal approaches its final choice (low *CV*_*end*_). We found that the *Lateral Displacement* set reached the highest score when following those parameters, suggesting that the transition from medial movement toward the screen to lateral movement toward the ports most effectively capture the mice’s choice.

#### Decision point uncertainty threshold selection

To estimate the timing of decision commitment, we defined a decision point as the final timepoint in each trial where the model’s predicted probability crossed a specific threshold. To capture the turning point, we identified quantitatively the threshold that best captures the behavioral transition from screen-approach to port-directed movements. This ensures that the decision point used in downstream analysis aligns with the animal’s physical change.

We evaluated five candidate uncertainty thresholds *T* ∈ {0.1, 0.2, 0.3, 0.4, 0.5}. For each threshold, we calculated the change (Δ) in four kinematic variables by comparing the mean values of the 10 timepoints immediately preceding and following the predicted decision point: x-velocity (expected increase), y-velocity (expected decrease), absolute head-to-body angle (expected increase), and running direction relative to the screen (expected increase). To account for inherent trial noise, each kinematic variable *k* was first Z-scored relative to a null distribution throughout the session, generated by randomly sampling two baseline windows within each trial:

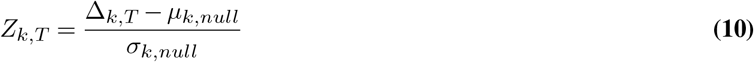

To ensure that variables contribute equally to the selection process, we applied min-max standardization across all candidate thresholds, so that each Z-scored variable is in [0, 1] while preserving the relative magnitude differences between thresholds:

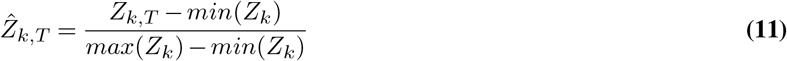

Finally, we calculated an aggregated turning score (*S*_*T*_) for each threshold:

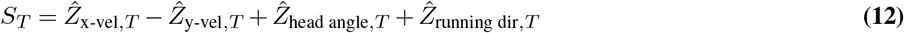

#### Decision points analysis

Once we selected the threshold that best captured the turning point in the trajectory (i.e. 0.2 uncertainty threshold, 80% confidence level), we compared the spatial coordinate of these decision points across occlusion conditions, and more precisely their distance to the screen.

To investigate a potential link between the task performance and the spatial location of the decision points, we compared the average success rate to the average decision point location per mouse. In order to capture the full range of performances for that specific analysis, we included all sessions that were above 70% success rate for more than 125 trials, but that did not pass the reward drop criterion. First, we fitted a linear least-squares regression across mice and reported the Pearson correlation coefficient and p-value (Wald test, t-distributed test statistic), using `scipy.stats.linregress`. We also identified the upper and lower quintile groups of sessions with decision points closest to or farthest from the screen under the narrow aperture condition and compared average success rates between these groups between mice (all sessions for each mouse, not only the sessions contained in the quintile groups). Because all mice performed above the criterion (success rate above 0.7), this analysis provides a stringent test for a potential influence of strategy on task execution.

## Supplemental Information Index

**Figures S1–S5, related to Figures 1–5**.

### Video S1. Dual-occlusion session example, related to Figure 3

Randomly sampled trials for an example session on the dual-occlusion task (session: Pheasant, 2024-08-15). Left: arena top view; Right: back view. Trials were extracted from trial onset, where virtual objects and occluders are spawned, to reward delivery; the inter-trial interval periods have been removed. Video playback speed: 0.3×.

### Video S2. Multi-occlusion session example, related to Figure 5

Randomly sampled trials for an example session on the multi-occlusion task (session: Nightingale, 2024-08-16). Left: arena top view; Right: back view. Trials were extracted from trial onset, where virtual objects and occluders are spawned, to reward delivery; the inter-trial interval periods have been removed. Video playback speed: 0.3×.

**Figure S1.**
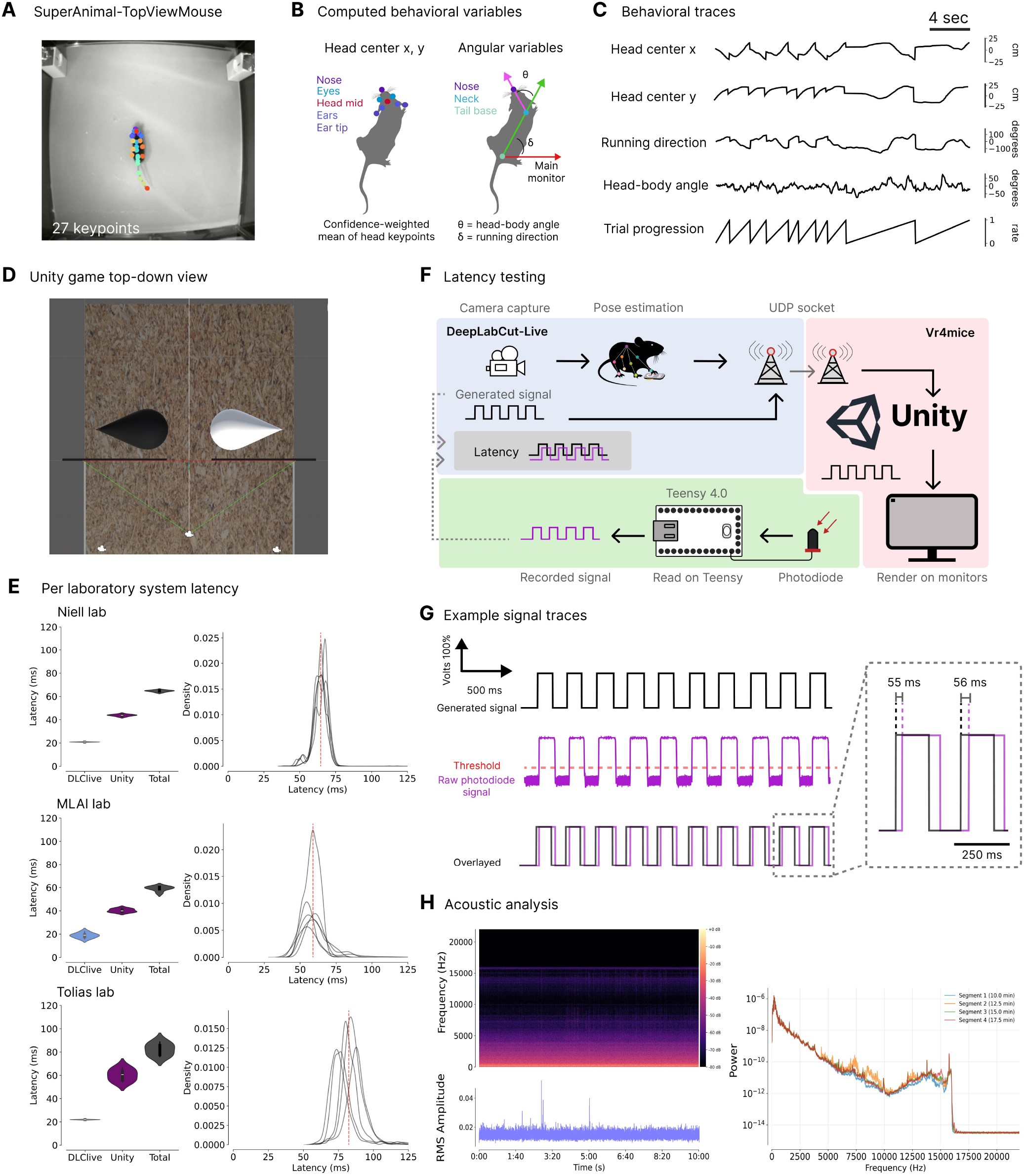
Architecture, latency profiling, and environmental validation of the virtual reality system, Related to Figure 1. **A**: Top view of a mouse in the virtual reality setup, with the corresponding 27 keypoints inferred using the SuperAnimal-TopViewMouse model. **B**: Computed behavioral variables fed to the game. Head center x and y position is calculated as the average of the head keypoints weighted by their confidence. The running direction relative to the front screen and the head angle relative to the body are also computed. **C**: Example traces of the behavioral variables and trial progression. **D**: Top-down view of the virtual arena in Unity during the aperture test. The red plane contains the 2 occluding walls, the camera in the center corresponds to the agent from which the viewpoint is rendered in the game. Here, the objects are visible through a narrow aperture. **E**: System latency per laboratory. Left: Camera-to-Render latency (total) and latencies of each processing step for respectively, the Niell laboratory (Up), MLAI laboratory (Middle) and Tolias laboratory (Down). The DeepLabCut-Live latency was computed as the time for the frame to be acquired and the pose inference and kinematic features to be calculated. The Unity latency corresponds to the time taken for the data to be sent to Unity and rendered on the screen. The camera-to-Render latency is the discrepancy in phase between the generated signal at frame capture and the recorded signal when the game is rendered on the monitor. Right: Individual latency profiles for individual latency tests. **F**: Setup architecture to test the latency of the system. A binary signal is generated in the DeepLabCut-Live processor, sent to the Unity game, rendered on the monitors and recorded by a photodiode via a Teensy 4.0. **G**: Example signal traces. In black is the signal generated in the DeepLabCut-Live processor, in purple is the signal recorded by the photodiode. A threshold is used to binarize the recorded signal and the total latency corresponds to the discrepancy between the generated signal and clean photodiode recordings. **H**: Acoustic stability analysis of the experimental environment on a 10-minute audio recordings during a session. (Top-Left): Long-term spectrogram of ambient noise. (Bottom-Left) Root Mean Square (RMS) of sound pressure levels. (Right): Segmented Power Spectral Density (PSD) comparison across four discrete 2.5-minute segments.

**Figure S2.**
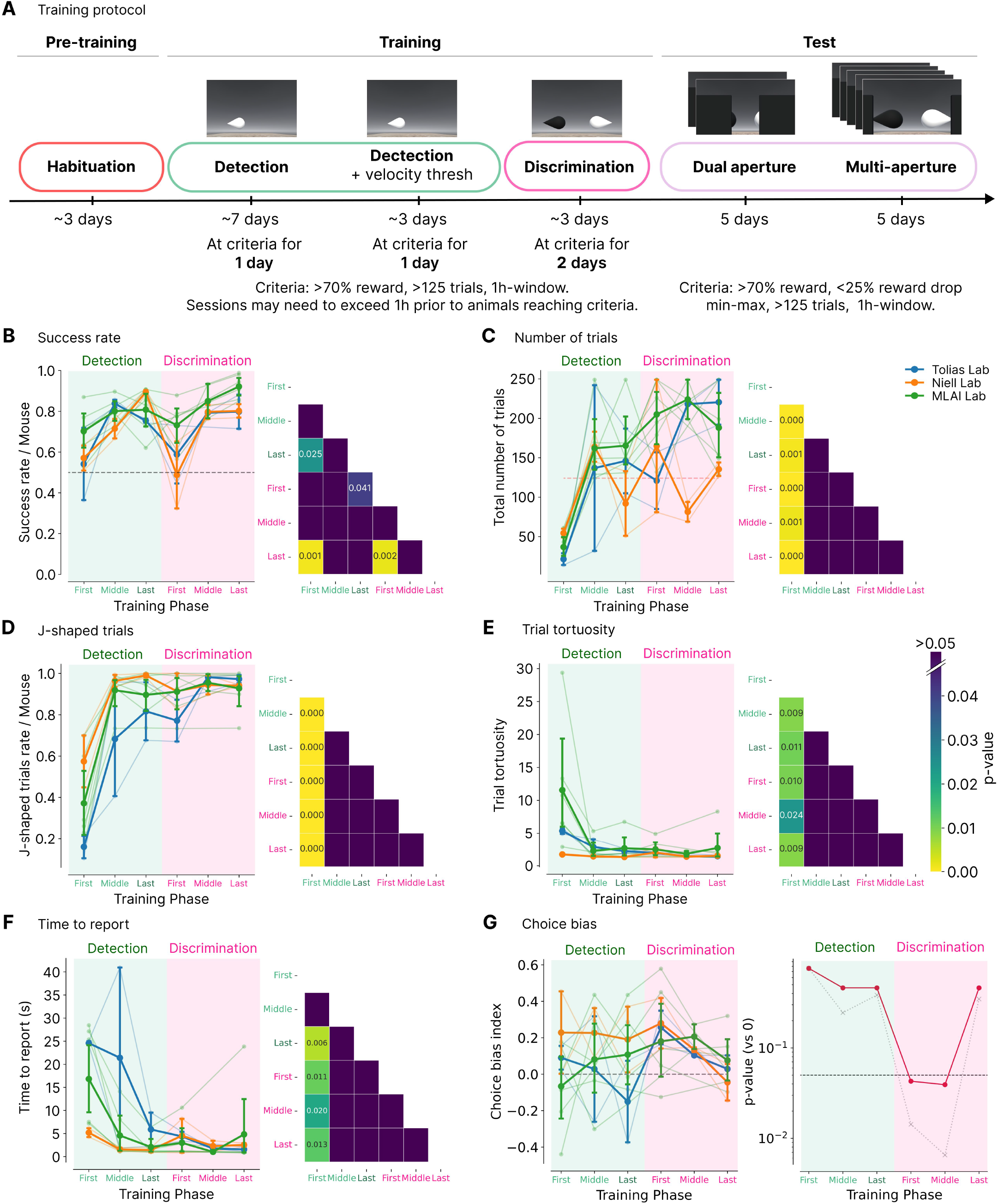
Training protocol, behavioral metrics and statistical comparisons across learning phases, Related to Figure 2. **A**: Overview of the experimental protocol followed in parallel in the three laboratories that ran the experiments. Left: In green, the detection phase with First and Middle sessions corresponding to the first and last sessions of the detection phase without velocity threshold and the Last session corresponding to the last session of the detection with velocity threshold. In pink, the First, Last and, if applicable, Middle sessions of the discrimination phase (with discriminator). Means ± s.e.m. (bold lines) per lab and individual sessions (thin lines) over the course of the training phases. Right: Except for **E**, corresponding heatmaps displaying FDR-corrected p-values for all pairwise comparisons following one-way ANOVA and post-hoc Tukey HSD tests. Significant differences are indicated through the colormap (*p <* 0.05), darker comparisons are non significant (*p >* 0.05). Legends are common to all panels. **B**: Success rate per mouse. One-way ANOVA significant effect of training stage, F(1, 53) = 8.13, *p* = 0.0062. **C**: Total number of trials. One-way ANOVA significant effect of training stage, F(1, 53) = 22.75, *p <* 0.0001. **D**: J-shaped trials rate. One-way ANOVA significant effect of training stage, F(1, 53) = 29.53, *p <* 0.0001. **E**: Trial tortuosity. One-way ANOVA significant effect of training stage, F(1, 53) = 8.33, *p* = 0.0056. **F**: Time to report in seconds. One-way ANOVA significant effect of training stage, F(1, 53) = 11.27, *p* = 0.0015. **G**: Choice bias index. Right: Log-scale, p-values (dashed) and FDR-corrected p-values (red) for one-sample t-tests comparing choice bias index against zero across training stages. First Detection: t = 0.32, *p* = 0.76; Middle Detection: t = 1.24, *p* = 0.25; Last Detection: t = 0.91, *p* = 0.38; First Discrimination: t = 3.03, *p* = 0.014; Middle discrimination: t = 5.20, *p* = 0.007; Last discrimination: t = 0.99, *p* = 0.35. Dashed line indicates significance threshold (*α* = 0.05). Only early and middle discrimination phases show significant bias away from zero.

**Figure S3.**
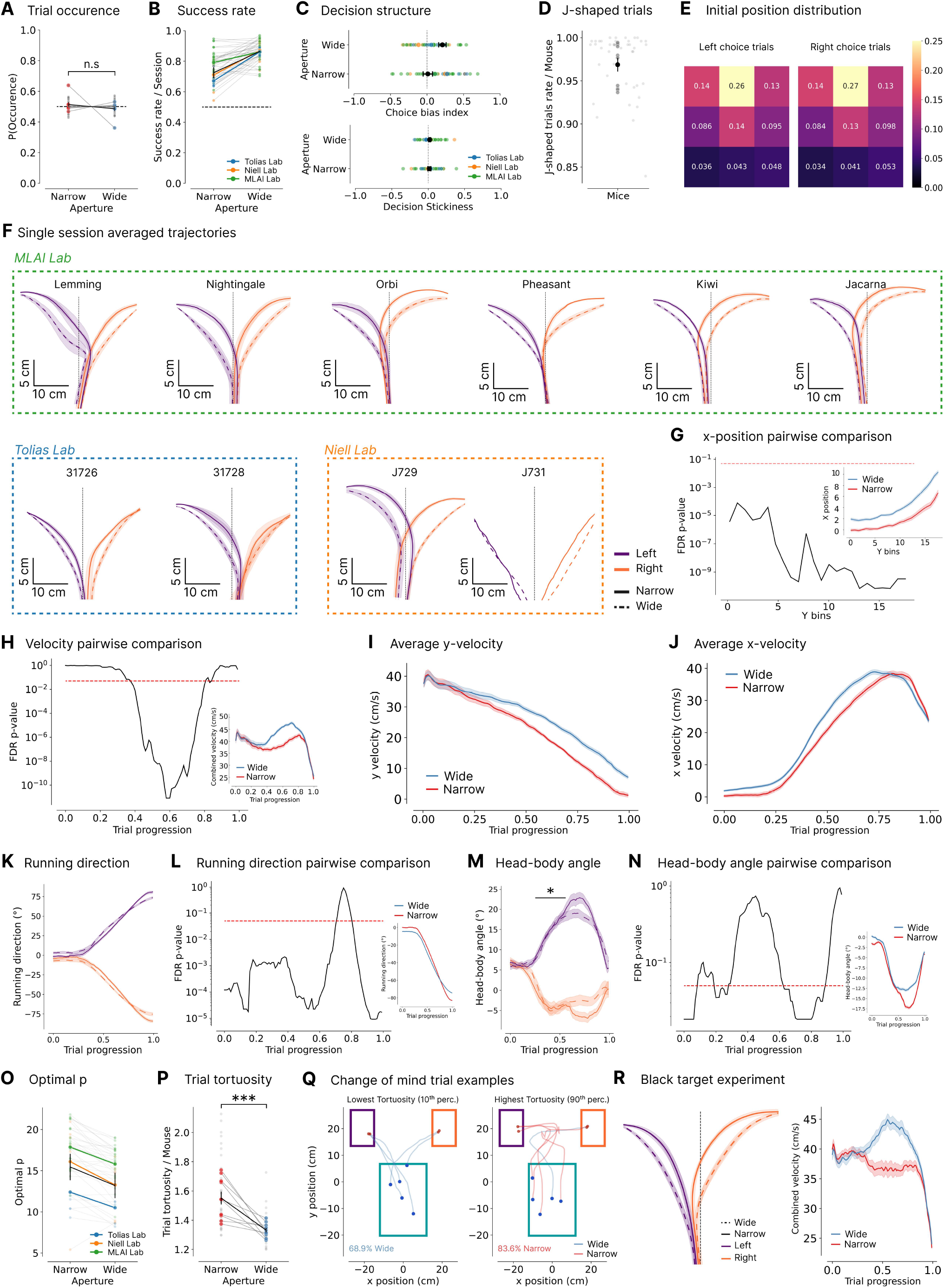
Dual-occlusion task: behavioral metrics, kinematic profiles, and trajectory analysis, Related to Figure 3. **A**: Relative trial proportions per aperture and mouse. n.s.: paired t-test: t(42) = 0.49, *p* = 0.63. Black: mean ± s.e.m.; colored: per-mouse; grey: per-session. **B**: Success rate by laboratory. **C**: Decision structure by laboratory. Above: side choice bias (0 = no bias, ± 1 = full left/right). Below: decision stickiness (0 = no stickiness, +1 = full stickiness, − 1 = full switching). **D**: J-shaped trials rate vs. wandering as engagement (threshold: duration ≤ 5 sec, tortuosity ≤ 5). Mean ± s.e.m. ± 0.973 0.005 (black); grey: per-mouse; small gray: per-session averages. **E**: Initial 3×3 position occupancy for the Left/Right choices. FDR-corrected paired t-test between choices per bin, t(43), *p >* 0.895 for all bins. **F**: Individual mouse mean trajectories ± s.e.m. (shaded) across sessions (Lemming: s=4; Nightingale: s=5; Oribi: s=5; Pheasant: s=5; Kiwi: s=5; Jacana: s=5; 31726: s=4; 31728: s=3; J729: s=6; J731: s=1). Dashed line: x-axis midline. Trajectories shown from arena midpoint to screen. **G**: Log-scale FDR-corrected paired t-tests on mean x-position per spatial bin between apertures (insert: interpolated position, flipped x, 0-18 cm in the arena). Two-way repeated-measures ANOVA: significant effects of aperture,: F(1, 42) = 68.65, spatial bin: F(40, 1680) = 89.21, and interaction: F(40, 1680) = 6.41; all *p* < 0.0001 (time-dependent differences in trajectory profile). **H**: Log-scale FDR-corrected paired t-tests on mean velocity per time bin between apertures (insert: velocity in cm/s vs. trial progression). Two-way repeated-measures ANOVA, significant effects of aperture: F(1, 42) = 58.36, time progression: F(99, 4158) = 48.18, and interaction: F(99, 4158) = 35.61; all *p* < 0.0001 (time-dependent differences in velocity profile). Mean ± s.e.m. across trial progression per apertures (n=43) for **I**: x-velocity (in cm/s), **J**: y-velocity (in cm/s), **K**: running direction relative to screen (in degrees, °). Paired t-test with multiple comparisons across trial progression, significant aperture effect (*p* < 0.05) across trial, except 70-80%. **L**: Log-scale FDR-corrected paired t-tests on mean running direction between apertures (insert: flipped running direction vs. trial progression). Two-way repeated-measures ANOVA, significant effects of aperture: F(1, 42) = 19.06, time bin: F(99, 4158) = 1344.43, and interaction: F(99, 4158) = 33.54; all *p* < 0.0001. **M**: Head angle relative to the body (°).∗ : FDR-corrected paired t-test with multiple comparisons across trial progression, significant aperture effect (*p* < 0.05) on the 60-90% portion of the trial. **N**: Log-scale FDR-corrected paired t-tests on mean head-body angle between apertures (insert: flipped head-body angle vs. trial progression). Two-way repeated-measures ANOVA, significant effects of aperture: F(1, 42) = 30.96, time progression: F(99, 4158) = 92.55, and interaction: F(99, 4158) = 5.83; all *p <* 0.0001. **O**: Optimal *p* of fitted L_*p*_ curves by laboratory. **P**: Trial tortuosity. Means ± s.e.m.: 1.56 ± 0.03 (narrow), 1.33 ± 0.01 (wide). Paired t-test: t(42) = −6.85, ∗ ∗ ∗: *p* = 2.41 × 10^*−*8^. **Q**: Example trials for 10^th^ and 90^th^ forward tortuosity percentiles, colored per aperture. Low tortuosity trials predominantly wide-aperture (68.9% wide vs. 31.1% narrow); high tortuosity trials primarily narrow-aperture (83.6% narrow vs. 16.4% wide). **R**: Black target results. Success rate; narrow: 0.74 ± 0.02, wide: 0.90 ± 0.02; t(14) = 4.35, *p* = 6.67 × 10^*−*4^. Left: Trajectory per aperture and choice. Two-way repeated-measures ANOVA, significant effects for aperture: F(1, 14) = 11.96, *p* = 0.0038, spatial bin: F(14, 196) = 221.97, *p* < 0.0001 and interaction: F(14, 196) = 5.80, *p <* 0.0001. FDR-corrected paired t-test with multiple comparisons across y bins, significant aperture effect (*p <* 0.05) from 2.5 cm of the initiation area top onward. Right: Combined velocity (m/s) per aperture. Two-way repeated-measures ANOVA, significant effect of aperture: F(1, 14) = 63.07, time bin: F(99, 1386) = 46.39, and interaction: F(99, 1386) = 22.22; all *p <* 0.0001. Paired t-test with multiple comparisons across the trial progression, significant aperture effect (*p <* 0.05) at 40-80% trial. Decision points; closer to the screen in the narrow-aperture trials; narrow: 14.64 ± 0.67, wide: 18.29 ± 0.52, mean difference: 3.65; t(14) = 3.76, *p* = 0.021). **B, C** and **O**: Mean ± s.e.m. (black), lab averages (colored), session averages (small colored).

**Figure S4.**
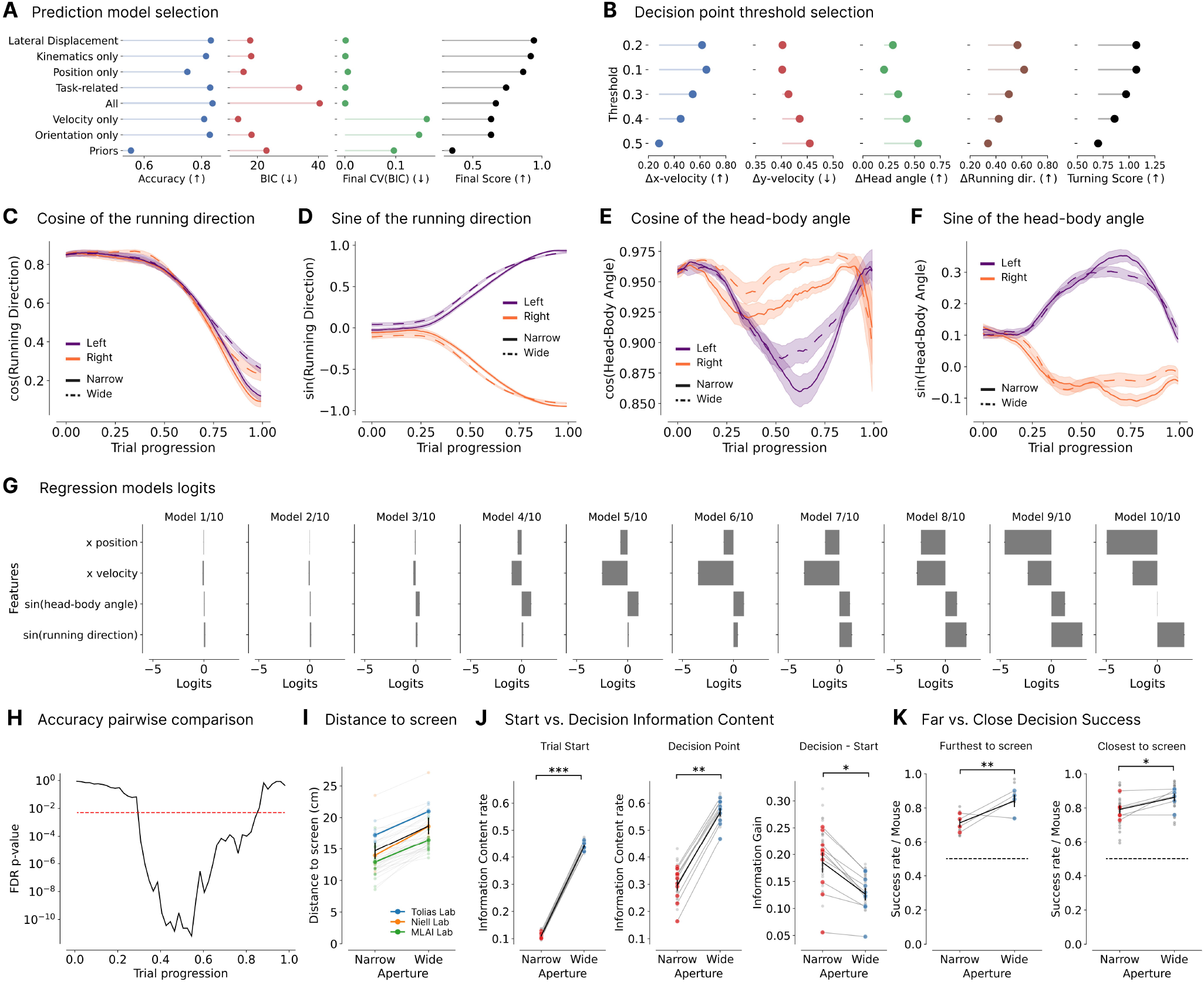
Dual-occlusion task: kinematic feature selection, decision variable regression, and spacial decision dynamics, Related to Figure 4. **A**: Feature set selection. Optimal set identified via three criteria: high accuracy, low BIC, and low BIC coefficient of variation for the final segment. The *Lateral Displacement* model was selected. **B**: Decision threshold selection. Turning point identified by evaluating uncertainty thresholds against shifts in four kinematic variables (x-velocity, y-velocity, head-body angle, and running direction) 10 timepoints pre/post-decision, aggregated into a “turning score”. Means ± s.e.m. across trial progression per aperture and choice for: **C**: cosine of the running direction, **D**: sine of the running direction, **E**: cosine of the head-body angle, and **F**: sine of the head-body angle. **G**: Logistic regression weights (logits) mean ± s.e.m. (n=43 sessions) for each segment model. **H**: Log-scale FDR-corrected paired t-tests on mean logistic regression accuracy between apertures (insert: accuracy vs. trial progression). Two-way repeated-measures ANOVA, significant effects of aperture: F(1, 42) = 26.19, time bin: F(49, 2058) = 794.25, and interaction: (F(49, 2058) = 9.63; all *p <* 0.0001. **I**: Decision point distance to screen (cm). Black: mean ± s.e.m.; colored: per-lab; thin colored: per session. **J**: Information content rate at trial onset (left) and decision point (middle), and their difference(right) per aperture. Paired t-test: ∗ ∗ ∗ : Onset: t(42) = 203.59, *p* = 1.58 × 10^*−*64^; ∗∗ : Decision, t(42) = 47.40, *p* = 4.26 × 10^*−*38^; ∗: Difference, t(42) = −10.97, *p* = 6.67 × 10^*−*14^. Black: mean ± s.e.m.; colored: per-lab; thin colored: per session. **K**: Success rate per aperture for groups with furthest (left) and closest (rigth) decisions points to screen. Individual session are in grey. Narrow vs. wide paired t-tests: furthest (4 mice, 13 sessions), ∗∗: paired t-test, t(12) = 8.62, *p* = 1.74 ×10^*−*6^.; closest (6 mice, 30 sessions), ∗: paired t-test, t(30) = 3.93, *p* = 4.81 ×10^*−*4^. Narrow-only groups comparison: t(41) = −3.16, *p* = 0.0029. Wide-only groups comparison: t(41) = 0.18, *p* = 0.86.

**Figure S5.**
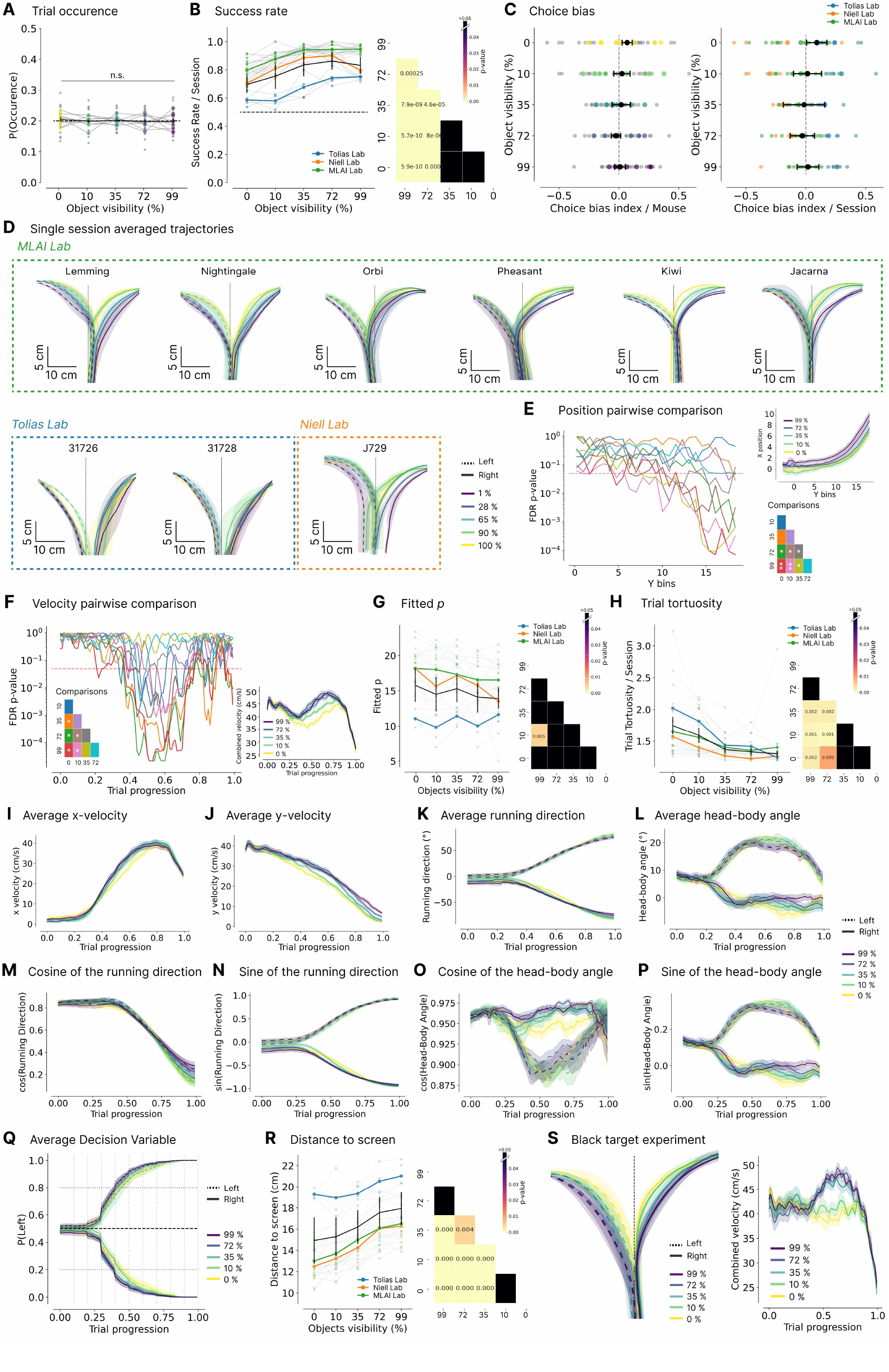
Multi-occlusion task: behavioral metrics, kinematic profiles, and decision variable modeling across visibility conditions, Related to Figure 5. **A**: Trial occurrence per aperture and mouse. n.s.: all paired t-tests *p* > 0.05. Black: mean ± s.e.m.; colored: per-mouse; grey: per-session. **B**: Success rate per aperture by laboratory. FDR-corrected paired t-tests: 99% and 72% visibility significantly differ from other conditions. Black: mean ± s.e.m.; colored: per-mouse; small colored: per-session. **C**: Side choice bias (0 = no bias, ±1 = full left/right) averaged per aperture for mice (Left) and laboratories (Right). Mean ± s.e.m.: 99%: 0.049 ± 0.034, 72%: –0.002 ± 0.037, 35%: –0.044 ± 0.029, 10%: –0.029 ±0.050, 0%: –0.015 ± 0.058. FDR-corrected t-tests vs. no bias: no significant deviation. 99%: t = 1.45, *p* = 0.160; 72%: t = –0.05, *p* = 0.964; 35%: t = –1.50, *p* = 0.147; 10%: t = –0.58, *p* = 0.568; 0%: t = –0.26, *p* = 0.798. **D**: Individual mouse trajectories ± s.e.m. (shaded) across sessions (Lemming: s=3; Nightingale: s=4; Oribi: s=3; Pheasant: s=2; Kiwi: s=3; Jacana: s=2; 31726: s=2; 31728: s=2; J729: s=4), grouped by laboratories. Dashed line: x-axis midline. Trajectories shown from arena midpoint to screen. Log-scale FDR-corrected paired t-tests between apertures for: **E**: Mean x-position per spatial bin (insert: flipped x, interpolated trials, 0-19 cm in the arena). Two-way repeated-measures ANOVA, significant effect of aperture: F(4, 96) = 8.08, Y position: F(28, 672) = 112.83, and interaction: F(112, 2688) = 3.45; all*p* < 0.0001. Paired t-test corrected p-value, ∗∗: *p* < 0.05 from zero onward, ∗ : *p <* 0.05 from 10.8 cm onward, **F**: Mean velocity per time bin (insert: velocity in cm/s on interpolated trials). Two-way repeated-measures ANOVA, significant effect of aperture: F(4, 96) = 14.17, time bin: F(99, 2376) = 41.89, and interaction: F(396, 9504) = 5.40; all *p* < 0.0001. Paired t-test corrected p-value, ∗ : *p* < 0.05 (40-80% of trial. **G**: Optimal *p* of fitted L_*p*_ curves. One-way repeated-measures ANOVA, significant aperture effect: F(4, 96) = 3.14, *p* = 0.018. FDR-corrected paired t-tests: only 0%-72% pair significant. **H**: Trial tortuosity. One-way repeated-measures ANOVA, significant effect of aperture, F(4, 96) = 11.85, *p <* 0.0001. FDR-corrected paired t-test: *p* < 0.009 for all except (72-99%), (10-35%), (0-35%), and (0-10%). Mean ± s.e.m. across trial progression per aperture for: **I**: x-velocity in cm/s, **J**: y-velocity in cm/s **K**: running direction in degrees (°), **L**: head-body angle in degrees (°), **M**: cosine of the running direction, **N**: sine of the running direction, **O**: cosine of the head-body angle, and **P**: sine of the head-body angle. **Q**: Mean ± s.e.m of the dynamic decision variable (logistic regression, *P* (*lef t*)) across time segments models and sessions per choice and aperture (n=25 sessions). **R**: Decision distance to screen (cm) per aperture and laboratories. FDR-corrected paired t-test: *p <* 0.004 for all but (0-10%) and (72-99%). Black: mean ± s.e.m.; colored: per-mouse; small colored: per-session. **S**: Black target results. Success rates: 0%: 0.741± 0.019, 10%: 0.825 ± 0.030, 35%: 0.906 ± 0.021, 72%: 0.961 ±0.016, 99%: 0.944± 0.016. Left: Averaged trajectory per aperture and choice conditions. Two-way repeated-measures ANOVA, significant effects for aperture, F(4, 36) = 4.25, *p* = 0.0064, and spatial bin, F(24, 216) = 135.89, *p* < 0.0001. Interaction between aperture and time bin was not significant, F(96, 864) = 1.03, *p* = 0.4042: while aperture size influenced the trajectories, this effect did not vary significantly across the spatial bins. Right: Averaged combined velocity (m/s) per aperture condition. Two-way repeated-measures ANOVA, significant effect of aperture: F(4, 36) = 8.02, time bin: F(99, 891) = 36.20, and interaction: F(396, 3564) = 4.93; all *p* < 0.0001. Decision points were closer to the screen in the narrower aperture trials: 0%: 13.785 ± 0.688, 10%: 14.765 ± 0.612, 35%: 16.407 ± 0.664, 72%: 17.583 ± 0.924, 99%: 17.385 ± 0.685.

## References

1. D. H. Hubel and T. N. Wiesel. Receptive fields of single neurones in the cat’s striate cortex. The Journal of Physiology, 148(3):574–591, October 1959. ISSN 0022-3751, 1469-7793. doi: 10.1113/jphysiol.1959.sp006308.

2. D. H. Hubel and T. N. Wiesel. Receptive fields, binocular interaction and functional architecture in the cat’s visual cortex. The Journal of Physiology, 160(1):106–154, January 1962. ISSN 0022-3751, 1469-7793. doi: 10.1113/jphysiol.1962.sp006837.

3. Wt Newsome and Eb Pare. A selective impairment of motion perception following lesions of the middle temporal visual area (MT). The Journal of Neuroscience, 8(6):2201–2211, June 1988. ISSN 0270-6474, 1529-2401. doi: 10.1523/JNEUROSCI.08-06-02201.1988.

4. Kh Britten, Mn Shadlen, Wt Newsome, and Ja Movshon. The analysis of visual motion: a comparison of neuronal and psychophysical performance. The Journal of Neuroscience, 12(12):4745–4765, December 1992. ISSN 0270-6474, 1529-2401. doi: 10.1523/JNEUROSCI.12-12-04745.1992.

5. Christopher P. Burgess, Armin Lak, Nicholas A. Steinmetz, Peter Zatka-Haas, Charu Bai Reddy, Elina A.K. Jacobs, Jennifer F. Linden, Joseph J. Paton, Adam Ranson, Sylvia Schröder, et al. High-Yield Methods for Accurate Two-Alternative Visual Psychophysics in Head-Fixed Mice. Cell Reports, 20(10):2513–2524, September 2017. ISSN 22111247. doi: 10.1016/j.celrep.2017.08.047.

6. Jackson J. Cone, Morgan L. Bade, Nicolas Y. Masse, Elizabeth A. Page, David J. Freedman, and John H.R. Maunsell. Mice Preferentially Use Increases in Cerebral Cortex Spiking to Detect Changes in Visual Stimuli. The Journal of Neuroscience, 40(41):7902–7920, October 2020. ISSN 0270-6474, 1529-2401. doi: 10.1523/JNEUROSCI.1124-20.2020.

7. Gautam Reddy, Venkatesh N. Murthy, and Massimo Vergassola. Olfactory Sensing and Navigation in Turbulent Environments. Annual Review of Condensed Matter Physics, 13(1):191–213, March 2022. ISSN 1947-5454, 1947-5462. doi: 10.1146/annurev-conmatphys-031720-032754.

8. Marcin Leszczynski and Charles E. Schroeder. The Role of Neuronal Oscillations in Visual Active Sensing. Frontiers in Integrative Neuroscience, 13:32, July 2019. ISSN 1662-5145. doi: 10.3389/fnint.2019.00032.

9. Alfred L. Yarbus. Eye Movements During Perception of Complex Objects. In Eye Movements and Vision, pages 171–211. Springer US, Boston, MA, 1967. ISBN 978-1-4899-5381-0 978-1-4899-5379-7. doi: 10.1007/978-1-4899-5379-7_8.

10. David Kleinfeld, Ehud Ahissar, and Mathew E Diamond. Active sensation: insights from the rodent vibrissa sensorimotor system. Current Opinion in Neurobiology, 16(4):435–444, August 2006. ISSN 09594388. doi: 10.1016/j.conb.2006.06.009.

11. Rolf J. Skyberg and Cristopher M. Niell. Natural visual behavior and active sensing in the mouse. Current Opinion in Neurobiology, 86:102882, June 2024. ISSN 09594388. doi: 10.1016/j.conb.2024.102882.

12. Scott Cheng-Hsin Yang, Máté Lengyel and Daniel M Wolpert. Active sensing in the categorization of visual patterns. eLife, 5:e12215, February 2016. ISSN 2050-084X. doi: 10.7554/eLife.12215.

13. Scott Cheng-Hsin Yang, Daniel M Wolpert, and Máté Lengyel. Theoretical perspectives on active sensing. Current Opinion in Behavioral Sciences, 11:100–108, October 2016. ISSN 23521546. doi: 10.1016/j.cobeha.2016.06.009.

14. James G. March. Exploration and Exploitation in Organizational Learning. Organization Science, 2(1):71–87, February 1991. ISSN 1047-7039, 1526-5455. doi: 10.1287/orsc.2.1.71.

15. Richard S. Sutton and Andrew G. Barto. Reinforcement learning: An introduction. MIT Press, 2018.

16. Eleanor M. Caves, Nicholas C. Brandley, and Sönke Johnsen. Visual Acuity and the Evolution of Signals. Trends in Ecology & Evolution, 33(5): 358–372, May 2018. ISSN 01695347. doi: 10.1016/j.tree.2018.03.001.

17. Mary Hayhoe and Dana Ballard. Eye movements in natural behavior. Trends in Cognitive Sciences, 9(4):188–194, April 2005. ISSN 13646613. doi: 10.1016/j.tics.2005.02.009.

18. Pedro Maldonado, Cecilia Babul, Wolf Singer, Eugenio Rodriguez, Denise Berger, and Sonja Grün. Synchronization of Neuronal Responses in Primary Visual Cortex of Monkeys Viewing Natural Images. Journal of Neurophysiology, 100(3):1523–1532, September 2008. ISSN 0022-3077, 1522-1598. doi: 10.1152/jn.00076.2008.

19. Amin Haji-Abolhassani and James J. Clark. An inverse Yarbus process: Predicting observers’ task from eye movement patterns. Vision Research, 103:127–142, October 2014. ISSN 00426989. doi: 10.1016/j.visres.2014.08.014.

20. A. Borji and L. Itti. Defending Yarbus: Eye movements reveal observers’ task. Journal of Vision, 14(3):29–29, March 2014. ISSN 1534-7362. doi: 10.1167/14.3.29.

21. Junji Ito, Cristian Joana, Yukako Yamane, Ichiro Fujita, Hiroshi Tamura, Pedro E. Maldonado, and Sonja Grün. Latency shortening with enhanced sparseness and responsiveness in V1 during active visual sensing. Scientific Reports, 12(1):6021, April 2022. ISSN 2045-2322. doi: 10.1038/s41598-022-09405-4.

22. Roland Baddeley, L. F. Abbott, Michael C. A. Booth, Frank Sengpiel, Tobe Freeman, Edward A. Wakeman, and Edmund T. Rolls. Responses of neurons in primary and inferior temporal visual cortices to natural scenes. Proceedings of the Royal Society of London. Series B: Biological Sciences, 264(1389):1775–1783, December 1997. ISSN 0962-8452, 1471-2954. doi: 10.1098/rspb.1997.0246.

23. Glen T Prusky, Paul W.R West, and Robert M Douglas. Behavioral assessment of visual acuity in mice and rats. Vision Research, 40(16): 2201–2209, July 2000. ISSN 00426989. doi: 10.1016/S0042-6989(00)00081-X.

24. Glen T. Prusky and Robert M. Douglas. Developmental plasticity of mouse visual acuity. European Journal of Neuroscience, 17(1):167–173, January 2003. ISSN 0953-816X, 1460-9568. doi: 10.1046/j.1460-9568.2003.02420.x.

25. Arne F. Meyer, John O’Keefe, and Jasper Poort. Two Distinct Types of Eye-Head Coupling in Freely Moving Mice. Current Biology, 30(11): 2116–2130.e6, June 2020. ISSN 09609822. doi: 10.1016/j.cub.2020.04.042.

26. Angie M Michaiel, Elliott Tt Abe, and Cristopher M Niell. Dynamics of gaze control during prey capture in freely moving mice. eLife, 9:e57458, July 2020. ISSN 2050-084X. doi: 10.7554/eLife.57458.

27. Philip Rl Parker, Elliott Tt Abe, Natalie T Beatie, Emmalyn Sp Leonard, Dylan M Martins, Shelby L Sharp, David G Wyrick, Luca Mazzucato, and Cristopher M Niell. Distance estimation from monocular cues in an ethological visuomotor task. eLife, 11:e74708, September 2022. ISSN 2050-084X. doi: 10.7554/eLife.74708.

28. Gioia De Franceschi, Tipok Vivattanasarn, Aman B. Saleem, and Samuel G. Solomon. Vision Guides Selection of Freeze or Flight Defense Strategies in Mice. Current Biology, 26(16):2150–2154, August 2016. ISSN 09609822. doi: 10.1016/j.cub.2016.06.006.

29. Samuel G. Solomon, Hadrien Janbon, Adam Bimson, and Thomas Wheatcroft. Visual spatial location influences selection of instinctive behaviours in mouse. Royal Society Open Science, 10(4):230034, April 2023. ISSN 2054-5703. doi: 10.1098/rsos.230034.

30. Jennifer L. Hoy, Iryna Yavorska, Michael Wehr, and Cristopher M. Niell. Vision Drives Accurate Approach Behavior during Prey Capture in Laboratory Mice. Current Biology, 26(22):3046–3052, November 2016. ISSN 09609822. doi: 10.1016/j.cub.2016.09.009.

31. Carl D Holmgren, Paul Stahr, Damian J Wallace, Kay-Michael Voit, Emily J Matheson, Juergen Sawinski, Giacomo Bassetto, and Jason Nd Kerr. Visual pursuit behavior in mice maintains the pursued prey on the retinal region with least optic flow. eLife, 10:e70838, October 2021. ISSN 2050-084X. doi: 10.7554/eLife.70838.

32. Aman B Saleem, Asli Ayaz, Kathryn J Jeffery, Kenneth D Harris, and Matteo Carandini. Integration of visual motion and locomotion in mouse visual cortex. Nature Neuroscience, 16(12):1864–1869, December 2013. ISSN 1097-6256, 1546-1726. doi: 10.1038/nn.3567.

33. Jinsung Chun, Youngsoo Kim, Jin Woo Choi, Daesoo Kim, and Sungho Jo. Egocentrically-stable discriminative stimulus-based spatial navigation in mice: implementation and comparison with allocentric cues. Scientific Reports, 9(1):6451, April 2019. ISSN 2045-2322. doi: 10.1038/s41598-019-42852-0.

34. Howard C. Boone, Jason M. Samonds, Emily C. Crouse, Carrie Barr, Nicholas J. Priebe, and Aaron W. McGee. Natural binocular depth discrimination behavior in mice explained by visual cortical activity. Current Biology, 31(10):2191–2198.e3, May 2021. ISSN 09609822. doi: 10.1016/j.cub.2021.02.031.

35. Cristopher M. Niell and Michael P. Stryker. Modulation of Visual Responses by Behavioral State in Mouse Visual Cortex. Neuron, 65(4):472–479, February 2010. ISSN 08966273. doi: 10.1016/j.neuron.2010.01.033.

36. Daniel A Dombeck, Christopher D Harvey, Lin Tian, Loren L Looger, and David W Tank. Functional imaging of hippocampal place cells at cellular resolution during virtual navigation. Nature Neuroscience, 13(11):1433–1440, November 2010. ISSN 1097-6256, 1546-1726. doi: 10.1038/nn.2648.

37. Guifen Chen, John Andrew King, Yi Lu, Francesca Cacucci, and Neil Burgess. Spatial cell firing during virtual navigation of open arenas by head-restrained mice. eLife, 7:e34789, June 2018. ISSN 2050-084X. doi: 10.7554/eLife.34789.

38. Eugene Balkovsky and Boris I. Shraiman. Olfactory search at high Reynolds number. Proceedings of the National Academy of Sciences, 99(20): 12589–12593, October 2002. ISSN 0027-8424, 1091-6490. doi: 10.1073/pnas.192393499.

39. Massimo Vergassola, Emmanuel Villermaux, and Boris I. Shraiman. ‘Infotaxis’ as a strategy for searching without gradients. Nature, 445(7126): 406–409, January 2007. ISSN 0028-0836, 1476-4687. doi: 10.1038/nature05464.

40. Jacqueline Gottlieb, Pierre-Yves Oudeyer, Manuel Lopes, and Adrien Baranes. Information-seeking, curiosity, and attention: computational and neural mechanisms. Trends in Cognitive Sciences, 17(11):585–593, November 2013. ISSN 13646613. doi: 10.1016/j.tics.2013.09.001.

41. Gary A Kane, Gonçalo Lopes, Jonny L Saunders, Alexander Mathis, and Mackenzie W Mathis. Real-time, low-latency closed-loop feedback using markerless posture tracking. eLife, 9:e61909, December 2020. ISSN 2050-084X. doi: 10.7554/eLife.61909.

42. Arbora Resulaj, Roozbeh Kiani, Daniel M. Wolpert, and Michael N. Shadlen. Changes of mind in decision-making. Nature, 461(7261):263–266, September 2009. ISSN 0028-0836, 1476-4687. doi: 10.1038/nature08275.

43. Diogo Peixoto, Jessica R. Verhein, Roozbeh Kiani, Jonathan C. Kao, Paul Nuyujukian, Chandramouli Chandrasekaran, Julian Brown, Sania Fong, Stephen I. Ryu, Krishna V. Shenoy, et al. Decoding and perturbing decision states in real time. Nature, 591(7851):604–609, March 2021. ISSN 0028-0836, 1476-4687. doi: 10.1038/s41586-020-03181-9.

44. Shih-Yi Tseng, Selmaan N. Chettih, Charlotte Arlt, Roberto Barroso-Luque, and Christopher D. Harvey. Shared and specialized coding across posterior cortical areas for dynamic navigation decisions. Neuron, 110(15):2484–2502.e16, August 2022. ISSN 08966273. doi: 10.1016/j.neuron.2022.05.012.

45. Alexander T. Lai, German Espinosa, Gabrielle E. Wink, Christopher F. Angeloni, Daniel A. Dombeck, and Malcolm A. MacIver. A robot-rodent interaction arena with adjustable spatial complexity for ethologically relevant behavioral studies. Cell Reports, 43(2):113671, February 2024. ISSN 22111247. doi: 10.1016/j.celrep.2023.113671.

46. David H. Gire, Vikrant Kapoor, Annie Arrighi-Allisan, Agnese Seminara, and Venkatesh N. Murthy. Mice Develop Efficient Strategies for Foraging and Navigation Using Complex Natural Stimuli. Current Biology, 26(10):1261–1273, May 2016. ISSN 09609822. doi: 10.1016/j.cub.2016.03.040.

47. John R Stowers, Maximilian Hofbauer, Renaud Bastien, Johannes Griessner, Peter Higgins, Sarfarazhussain Farooqui, Ruth M Fischer, Karin Nowikovsky, Wulf Haubensak, Iain D Couzin, et al. Virtual reality for freely moving animals. Nature Methods, 14(10):995–1002, October 2017. ISSN 1548-7091, 1548-7105. doi: 10.1038/nmeth.4399.

48. Manu S. Madhav, Ravikrishnan P. Jayakumar, Shahin G. Lashkari, Francesco Savelli, Hugh T. Blair, James J. Knierim, and Noah J. Cowan. The Dome: A virtual reality apparatus for freely locomoting rodents. Journal of Neuroscience Methods, 368:109336, February 2022. ISSN 01650270. doi: 10.1016/j.jneumeth.2021.109336.

49. Domonkos Pinke, John B. Issa, Gabriel A. Dara, Gergely Dobos, and Daniel A. Dombeck. Full field-of-view virtual reality goggles for mice. Neuron, 111(24):3941–3952.e6, December 2023. ISSN 08966273. doi: 10.1016/j.neuron.2023.11.019.

50. Linda Judák, Gergely Dobos, Katalin Ócsai, Eszter Báthory, Huba Szebik, Balázs Tarján, Pál Maák, Zoltán Szadai, István Takács, Balázs Chiovini, et al. Moculus: an immersive virtual reality system for mice incorporating stereo vision. Nature Methods, 22(2):386–398, February 2025. ISSN 1548-7091, 1548-7105. doi: 10.1038/s41592-024-02554-6.

51. Kathleen E. Cullen. Vestibular processing during natural self-motion: implications for perception and action. Nature Reviews Neuroscience, 20(6): 346–363, June 2019. ISSN 1471-003X, 1471-0048. doi: 10.1038/s41583-019-0153-1.

52. Eero P Simoncelli and Bruno A Olshausen. Natural Image Statistics and Neural Representation. Annual Review of Neuroscience, 24(1):1193– 1216, March 2001. ISSN 0147-006X, 1545-4126. doi: 10.1146/annurev.neuro.24.1.1193.

53. James J. DiCarlo, Davide Zoccolan, and Nicole C. Rust. How Does the Brain Solve Visual Object Recognition? Neuron, 73(3):415–434, February 2012. ISSN 08966273. doi: 10.1016/j.neuron.2012.01.010.

54. Samuel J. Gershman, Eric J. Horvitz, and Joshua B. Tenenbaum. Computational rationality: A converging paradigm for intelligence in brains, minds, and machines. Science, 349(6245):273–278, July 2015. ISSN 0036-8075, 1095-9203. doi: 10.1126/science.aac6076.

55. Herbert A. Simon. Theories of bounded rationality. In C. B. McGuire and R. Radner, editors, Decision and Organization, pages 161–176. lsevier, 1972.

56. Zhengwei Wu, Minhae Kwon, Saurabh Daptardar, Paul Schrater, and Xaq Pitkow. Rational thoughts in neural codes. Proceedings of the National Academy of Sciences, 117(47):29311–29320, November 2020. ISSN 0027-8424, 1091-6490. doi: 10.1073/pnas.1912336117.

57. Python Software Foundation. Python Language Reference, version 3.9–3.10, 2020.

58. Dimitri Yatsenko, Jacob Reimer, Alexander S. Ecker, Edgar Y. Walker, Fabian Sinz, Philipp Berens, Andreas Hoenselaar, R. James Cotton, Athanassios S. Siapas, and Andreas S. Tolias. DataJoint: managing big scientific data using MATLAB or Python, November 2015.

59. Alexander Mathis, Pranav Mamidanna, Kevin M. Cury, Taiga Abe, Venkatesh N. Murthy, Mackenzie Weygandt Mathis, and Matthias Bethge. DeepLabCut: markerless pose estimation of user-defined body parts with deep learning. Nature Neuroscience, 21(9):1281–1289, September 2018. ISSN 1097-6256, 1546-1726. doi: 10.1038/s41593-018-0209-y.

60. Tanmay Nath, Alexander Mathis, An Chi Chen, Amir Patel, Matthias Bethge, and Mackenzie Weygandt Mathis. Using DeepLabCut for 3D markerless pose estimation across species and behaviors. Nature Protocols, 14(7):2152–2176, July 2019. ISSN 1754-2189, 1750-2799. doi: 10.1038/s41596-019-0176-0.

61. Shaokai Ye, Anastasiia Filippova, Jessy Lauer, Steffen Schneider, Maxime Vidal, Tian Qiu, Alexander Mathis, and Mackenzie Weygandt Mathis. SuperAnimal pretrained pose estimation models for behavioral analysis. Nature Communications, 15(1):5165, June 2024. ISSN 2041-1723. doi: 10.1038/s41467-024-48792-2.

62. Géry Casiez, Nicolas Roussel, and Daniel Vogel. 1 C filter: a simple speed-based low-pass filter for noisy input in interactive systems. In Proceedings of the SIGCHI Conference on Human Factors in Computing Systems, pages 2527–2530, Austin Texas USA, May 2012. ACM. ISBN 978-1-4503-1015-4. doi: 10.1145/2207676.2208639.

63. Charles R. Harris, K. Jarrod Millman, Stéfan J. Van Der Walt, Ralf Gommers, Pauli Virtanen, David Cournapeau, Eric Wieser, Julian Taylor, Sebastian Berg, Nathaniel J. Smith, et al. Array programming with NumPy. Nature, 585(7825):357–362, September 2020. ISSN 0028-0836, 1476-4687. doi: 10.1038/s41586-020-2649-2.

64. Arthur Juliani, Vincent-Pierre Berges, Ervin Teng, Andrew Cohen,Jonathan Harper, Chris Elion, Chris Goy, Yuan Gao, Hunter Henry, Marwan Mattar, et al. Unity: A General Platform for Intelligent Agents, 2018. Version Number: 2.

65. Blender Online Community. Blender, 2024. Version Number: 4.1.

66. Carolina Cruz-Neira, Daniel J. Sandin, Thomas A. DeFanti, Robert V. Kenyon, and John C. Hart. The CAVE: audio visual experience automatic virtual environment. Communications of the ACM, 35(6):64–72, June 1992. ISSN 0001-0782, 1557-7317. doi: 10.1145/129888.129892.

67. Pauli Virtanen, Ralf Gommers, Travis E. Oliphant, Matt Haberland, Tyler Reddy, David Cournapeau, Evgeni Burovski, Pearu Peterson, Warren Weckesser, Jonathan Bright, et al. SciPy 1.0: fundamental algorithms for scientific computing in Python. Nature Methods, 17(3):261–272, March 2020. ISSN 1548-7091, 1548-7105. doi: 10.1038/s41592-019-0686-2.

68. Jeffrey S. Taube. Head direction cells and the neurophysiological basis for a sense of direction. Progress in Neurobiology, 55(3):225–256, June 1998. ISSN 03010082. doi: 10.1016/S0301-0082(98)00004-5.

69. Lutfiyya N. Muhammad. Guidelines for repeated measures statistical analysis approaches with basic science research considerations. Journal of Clinical Investigation, 133(11):e171058, June 2023. ISSN 1558-8238. doi: 10.1172/JCI171058.

70. Yoav Benjamini and Yosef Hochberg. Controlling the False Discovery Rate: A Practical and Powerful Approach to Multiple Testing. Journal of the Royal Statistical Society Series B: Statistical Methodology, 57(1):289–300, January 1995. ISSN 1369-7412, 1467-9868. doi: 10.1111/j.2517-6161.1995.tb02031.x.

71. Fabian Pedregosa, Gaël Varoquaux, Alexandre Gramfort, Vincent Michel, Bertrand Thirion, Olivier Grisel, Mathieu Blondel, Andreas Müller, Joel Nothman, Gilles Louppe, et al. Scikit-learn: Machine Learning in Python, 2012. Version Number: 4.

72. Gideon Schwarz. Estimating the Dimension of a Model. The Annals of Statistics, 6(2), March 1978. ISSN 0090-5364. doi: 10.1214/aos/1176344136.

73. Ernst Wit, Edwin Van Den Heuvel, and Jan-Willem Romeijn. ‘All models are wrong…’: an introduction to model uncertainty. Statistica Neerlandica, 66(3):217–236, August 2012. ISSN 0039-0402, 1467-9574. doi: 10.1111/j.1467-9574.2012.00530.x.

